# Specialized terminology limits the reach of new scientific knowledge

**DOI:** 10.1101/2020.08.20.258996

**Authors:** Alejandro Martínez, Stefano Mammola

## Abstract

Words are the building blocks of science. As our understanding of the world progresses, scientific disciplines naturally enrich their specialized vocabulary (jargon). However, in the era of interdisciplinarity, the use of jargon may hinder effective communication amongst scientists that do not share a common linguistic background. The question of how jargon limits the transmission of scientific knowledge has long been debated, but rarely addressed quantitatively. We explored the relationship between the use of jargon and citations using 21,486 articles focusing on cave research, a multidisciplinary field particularly prone to terminological specialization and where linguistic disagreement among peers is frequent. We demonstrate that the use of jargon in the title and abstract significantly reduces the number of citations a paper receives. Given that these elements are the hook to readers, we urge scientists to restrict jargon to sections of the paper where its use is unavoidable.

## MAIN TEXT

A stumbling toddler babbling “mummy” or a famous scholar writing his 500-pages lifetime essay have at least one thing in common: they both navigate reality through words. We all do, as much as we can speak or read. “*The limits of our words are the limits of our world*”, believed philosopher Ludwig Wittgenstein (1933) and not surprisingly, our education is largely devoted to learn new terms and their meanings. Whether it is a zoologist defining a white blind salamander as a “*neotenic metazoans with anophthalmia*”, or a geologist describing marble as a “*metamorphic rock produced by the recrystallization of calcite or dolomite”*, the importance of specialized terminology (jargon) is undisputed. Jargon, although difficult at first, condense years of knowledge into a precise mental image: *metazoan* depicts a *multicellular eukaryote*; hence an organism consisting of multiple cells with a *nucleus*; which brings up, if we understand this jargon, images of membranous structures containing the salamander’s genetic information. As in a matryoshka, each new term enriches the initial message with information, structuring and systematizing concepts into the corpus of Science (Hoyningen-Huene 2013)

However, words are not Science. Physicist Richard Feynman (1966) believed that learning the meaning of words will only inform about the limit of people’s imagination, but nothing about nature itself. Jargon illustrates complex concepts only in the minds of those sharing a common background, while precluding everyone else from understanding. When stepping out of its linguistic comfort zone, a reader might not understand the jargon at all, get the message only partially or, after an extra mental effort, figure it out its meaning. In other words, upon the *neotenic metazoan* above, a reader might be able to picture an Olm, imagine some-sort-of-weird-animal, or throw up his arms in despair upon a complete enigma. It is within this range of confusion where the evil of jargon abuse expresses himself in all its glory: not only it reinforces the distinction between a geologist and a zoologist but potentially, it also divides zoologists into an endless number of subgroups. In conclusion, jargon, as certain type of jokes, may communicate ideas powerfully (Žižek 2014), but also and perhaps more often, artificially define “insiders” and “outsiders”, reinforcing the isolation of academics within their respective ivory towers. And, as the abuse of jokes, too much jargon can be tiring (Pennisi 2016)

This is nothing new. Again and again, scientists (Montgomery 1989; Adams et al. 1997; Hirst 2003; Rakedzon et al. 2017; Barnett & Doubleday 2020), economists (Tan et al. 2019), and philosophers (Wittgenstein 1953) have warned us about the dangers of jargon abuse. *“Never use a […] jargon word if you can think of an everyday English equivalent”*, journalists George Orwell (1968) famously stated. However, as important as these scholars might be, their opinions remain subjective as long as the effect of jargon (ab)use on scientific reach remains unquantified. Recently, Plavén-Sigray et al. (2017) analyzed the abstract of >700,000 articles across 12 sub-disciplines of life and medical sciences, showing that an increase in the use of jargon decreases the readability of texts. However, this study focused on general scientific jargon such as “robust”, “therefore”, and “underlying”. This *robust* analysis *therefore underlies* the need of using a plain language when communicating with the public. Conversely, since these terms are now integrant part of scientist’s writing routine, it is unlikely that they will undermine communication amongst them. What remains to be quantified is the role of discipline-specific jargon in driving the impact of a paper across scientists with different backgrounds. Such an analysis could hardly be performed on a broad multidisciplinary database such as the one used by Plavén-Sigray et al. (2017), because the diversity of specialized terms and the factors affecting their use vary too much across disciplines. These two confounding factors, however, can be alleviated by looking at the literature within a single multidisciplinary community of scientists. In such a context, jargony papers will be less understood, remembered, and ultimately cited.

Since the early 20^th^ century (Racovitza 1907), cave research has been the colliding point of four generations of scientists with diverse scientific backgrounds. Geologists, zoologists, anthropologists, ecologists, and evolutionary biologists have promiscuously interacted in the darkness of caves populating 120 years of cave literature with a maze of specialized terms, either borrowed from their scientific backgrounds or just coined *ex novo* using diverse etymological roots (Appendix S1). The lack of terminological agreement amongst cave scientists have preserved most of these words, which are still commonly found in the literature. Even today, in the 21^st^ century, several of these terms are the central subject of heated etymological debates (Figure 1). We took advantage of the long tradition of multidisciplinary (Poulson and White 1969) and high terminological specialization (Culver and Pipan 2018) offered by cave literature, to investigate the effect of jargon use on article success—measured as the number of citations.

**Figure 1.**
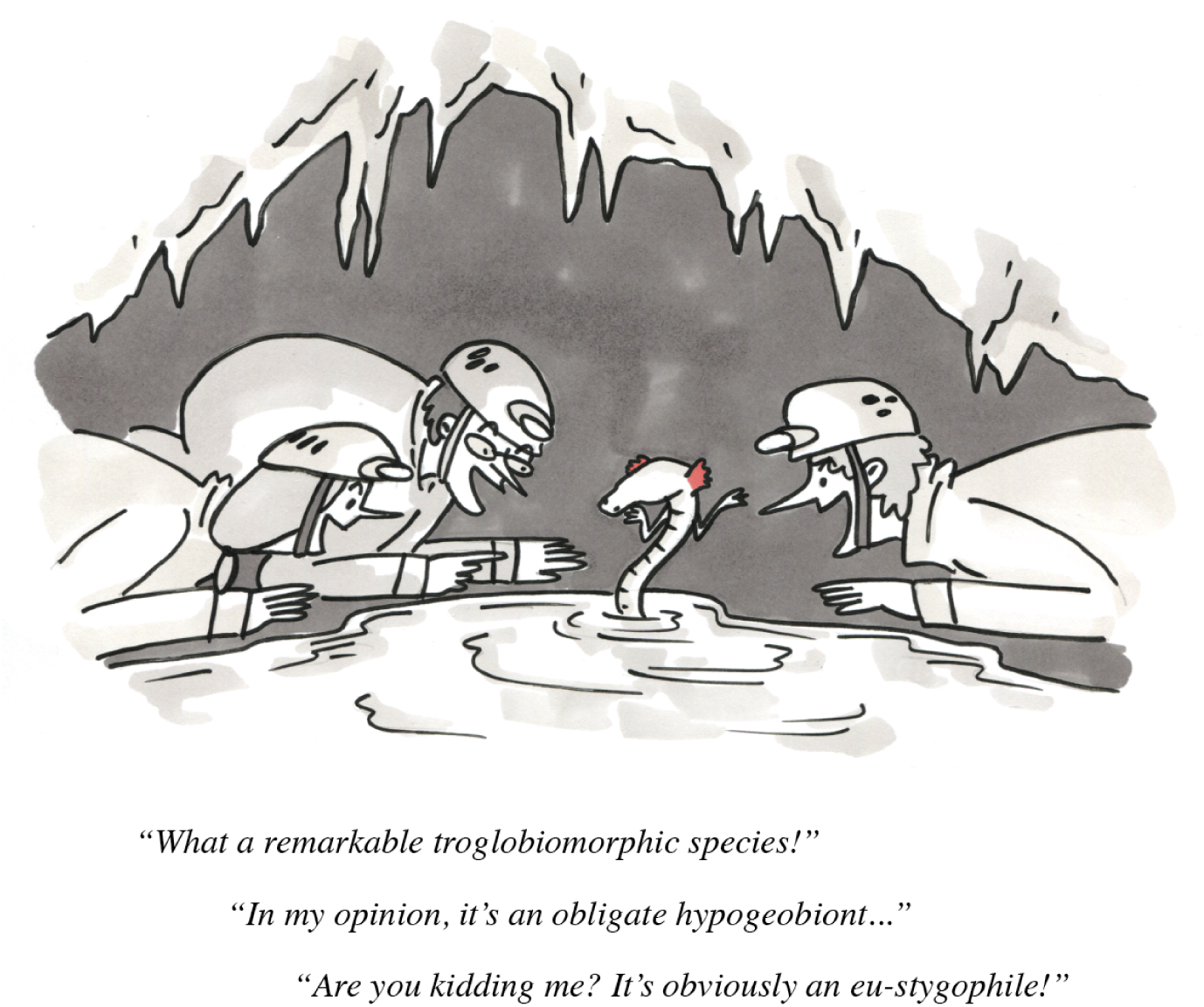
Jargon underground. Another day of terminological debate in the cave-office. Since the 19^th^ century, much ink has been spilled to discuss terminological nuances regarding the ecological classification of the subterranean fauna (see, e.g., Sket 2008; Giachino and Vailati 2017; Trajano and de Carvalho 2017), even though such classifications are just an attempt to simplify nature complexity, which is far from being rigidly defined (Mammola 2019). Illustration by Irene Frigo (https://www.instagram.com/irene.frigo/).

In the Web of Science, we sourced 21,486 research articles on cave environments published over the last 30 years. By using a curated selection of keywords, we ensured to cover articles dealing with caves published in both cave-specific and general international journals. To define discipline-specific jargon, we manually assembled a comprehensive list of c. 1500 words using glossaries of books focused on caves, reviews, and other sources (full list in Appendix S1). We calculated the proportion of jargon in title and abstract of each article over the total number of words. We focused on titles and abstracts given that these elements are the hook to readers (Mabe and Amin 2002) and reflect the overall writing style of entire articles (Plavén-Sigray et al. 2017). A detailed description of the analysis is in Appendix S2.

We observed a negative and non-linear effect of jargon on the number of citations, which significantly decreased as the proportion of jargon in the title (Figure 2a) and abstract increased (Figure 2c). This trend was particularly evident in abstracts, with a sudden drop in citations when the proportion of jargon was above 1% (Figure 2c inset). Interestingly, none of the highly cited papers (Pearson’s residuals of citations >100, corresponding to >450 citations) used jargon in the title, and almost all highly cited papers had a proportion of jargon in the abstract below 1%. All in all, the type of specialized words used in abstract and title were similar, although occurring with different frequencies (Figure 2b, d). We also found a positive correlation between the use of jargon in the title and abstract, with only about one third of articles using jargon in the abstract also including jargon in the title (Figure 3).

**Figure 2.**
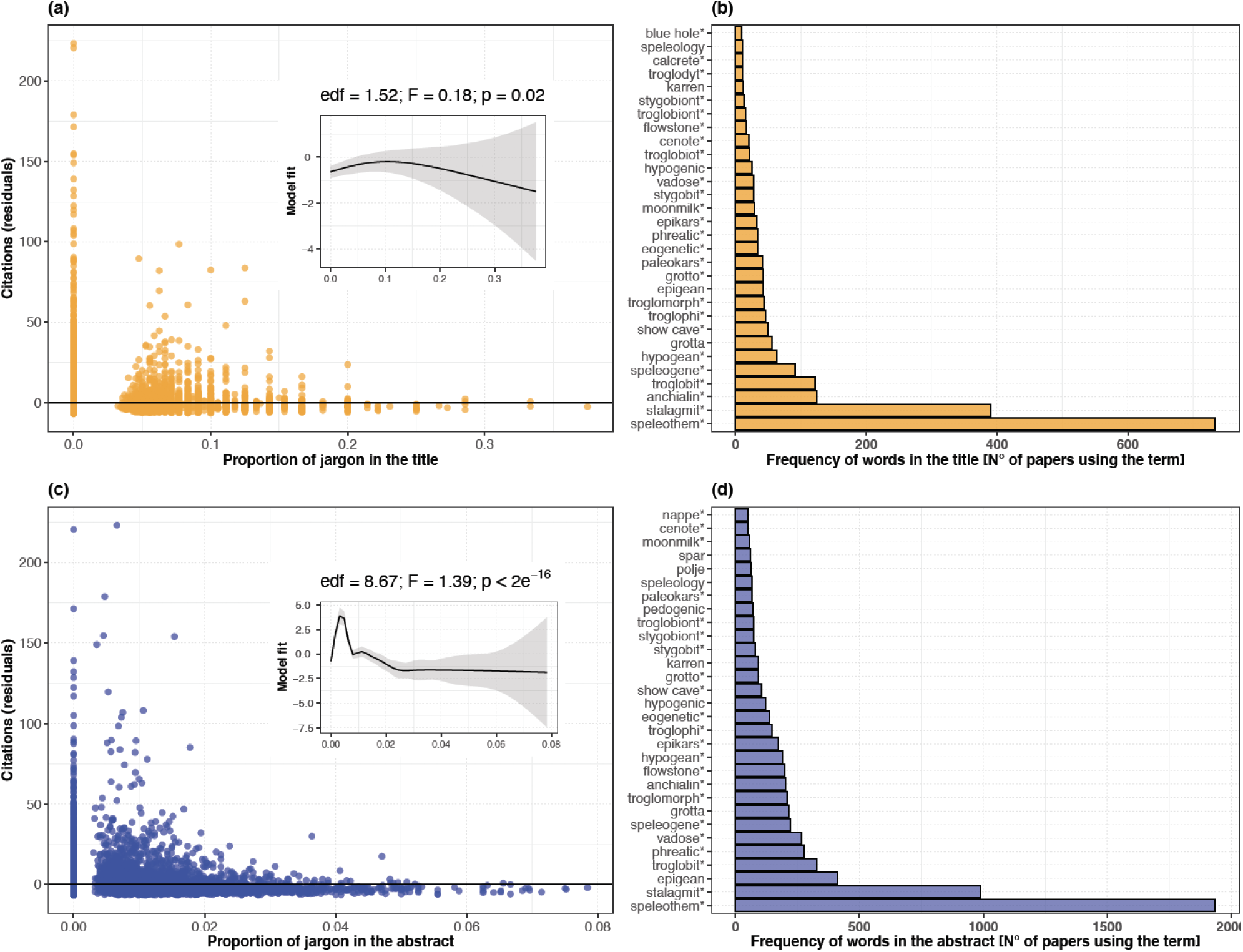
Effect of jargon use on citations and most frequently used jargon. **a, c)** Relationship between proportion of jargon and citations. Number of citations for each article is normalized by its age, expressing it as the Pearson’s residuals from the regression curve (Figure S1) representing the predicted number of citations over time (Mammola et al. 2020). Dots below the horizontal black line are articles under-cited for their age, and *vice versa*. Insets show the predicted trend based on Generalized additive mixed models, with a random structure to account for similarity of jargon between articles published in the same subject area and the variation of jargon through time. **b, d**) 30 most frequently used jargon terms in titles and abstracts. Using the regular expression notation, the asterisk (*) at the end of the words is a metacharacter for zero or more instances of the preceding characters (e.g., “speleogenom*” matches “speleogenome”, “speleogenomics”, etc.).

**Figure 3.**
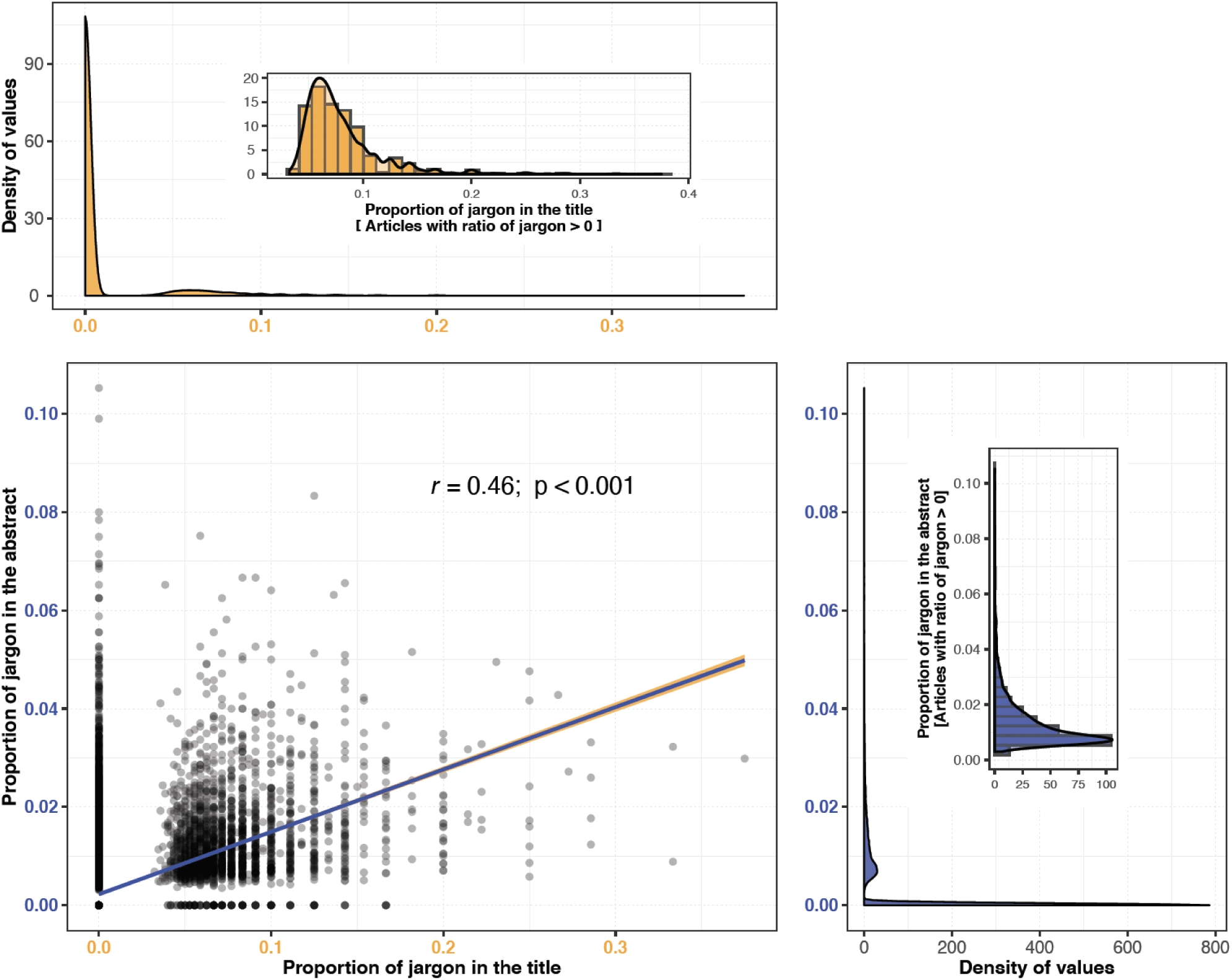
Correlation between the proportion of jargon in the abstract and title. Correlation is based the Pearson’s *r*. Density plots show the distribution of proportion of jargon values, obtained by computing a kernel density estimate. Insets are distribution of proportion of jargon values only for those articles with abstract or titles including jargon (i.e., proportion > 0).

While our analysis does not inform about the epistemological basis driving the choice for one word or another, it clearly emphasizes the negative effect of jargon on the success of a paper. With an estimated 1.5 million new articles published every year (Laurance et al. 2013), there is increasing pressure to publish papers that stand out amidst so many others (França and Montserrat 2019). A global estimate pointed out that scientists skim an average of over 1,100 titles and 200 abstracts a year, but they go on reading 97 full-texts (Mabe and Amin 2002). This suggests that the stylistic features of titles and abstracts act as important filters (Bowman and Kinnan 2018; França and Montserrat 2019; Freeling et al. 2019): if overuse of jargon prevents a reader from understanding the message of a paper, this paper is unlikely to end up being amongst the 97 chosen few. Given that the title and abstract bait readers interest, scientists might want to restrict jargon use to sections of the paper where its use is unavoidable.

In order to raise awareness to this problem, we would like to conclude by introducing a new jargon: the “Wittgensteinian Shortfall”. This obscure yet sophisticated combination of terms associates the philosophical ideas of the late Wittgenstein (1958) with the shortfall metaphor widely used in ecology (Hortal et al. 2015). While we might define it as “*the discrepancy between widely accessible scientific facts and the complexity of words used to name them”*, the real aim of the Wittgensteinian Shortfall is to create a “secret language” that we want to share exclusively with our readers. Our intent is to make apparent the wisdom of our elitarian community (Sand-Jensen 2007), allowing those who have read this paper to engage on the same *word games*. We believe that even jargonists will appreciate the irony in this.

## Acknowledgements

This paper started as a discussion with our friend Martina Pavlek in the Simplon Music Café in Pallanza. By reading with hindsight some of our early manuscripts, we realized how rich in jargon—and, in parallel, poorly cited—they were: this further stimulated us in exploring this subject. We thank Irene Frigo for preparing the illustrations, and Irene and Maria Bogolomova for helping us to avoid jargon in our everyday lives.

## Author contribution statement

AM sourced articles in Web of Science, curated the list of jargon words, and prepared the glossary. SM performed analyses and prepared figures. Both authors equally contributed to the writing.

## Data availability

Data and script to generate the analyses will be deposited in a public repository upon acceptance in a peer-reviewed journal.

## Supplementary material

**Appendix S1** – List of jargon terms used for the analyses.

**Appendix S2** – Extended methods.

## Notes

### Competing Interest Statement

The authors have declared no competing interest.

