## Appendix S2 for "Specialized terminology limits the reach of new scientific knowledge"

### EXTENDED METHODS

#### 1. Selection of articles

We aimed to obtain a representative set of articles focused on cave and other subterranean environments (e.g., groundwaters) published in international scientific journals. We restricted this analysis to cave-based science because it is a discipline particularly rich in specialized terminology (see Figure 1 and discussion in the main text). Moreover, the study of subterranean habitats is essentially an interdisciplinary endeavors in that it requires some understanding in diverse research areas, from ecology and biology, to geology and paleontology, to climatology and physics, and even human cultural evolution (Moldovan et al., 2018; Culver & Pipan, 2019).

We sourced articles in the Web of Science (Clarivate Analytics), a database containing metadata of scientific literature items published between 1965 and 2019. We performed the search on 1st March 2020 in Pallanza (Italy), using the browser Google Chrome (Pozsgai et al. 2020). We considered only articles written in English. We used a searching string designed to capture articles in all research areas potentially encompassing cave-based studies (geology, subterranean biology, paleontology, archeology, etc.), as follows:

TS = ("cave") OR TS = ("caves") OR TS = ("subterranean biolog *") OR TS = ("biospel *") OR TS = ("speleo *") OR SO = ("Subterranean biology") OR SO = ("Acta Carsologica") OR SO = ("International Journal of Speleology") OR SO = ("Journal of Cave and Karst Studies") OR SO = ("Stygologia")

The string is in the Web of Science notation, where:

i) TS indicate searches for general ‘topics’;

ii) SO indicate searches for specific journal names;

iii) and the asterisk (*) is a regular expression indicating to match all words including that string of characters (for example, “biospel*” will match “biospeleology”, biospeleologists”, and “biospeleological”).

The inclusion of search terms for specific journals (SO) was designed to ensure capturing all articles published in the few specialized journals entirely devoted to publishing research in subterranean environments. The initial search yielded 24,569 published items, encompassing different research areas (6,418 in Geoscience multidisciplinary; 2,541 Archeology; 2,846 Geography physical; 2,526 Zoology; 2,233 Anthropology; 1,364 Multidisciplinary science; 1,306 Ecology; 1,088 Evolutionary biology; 1,070 Geology; 1,031 Environmental science; and 3177 Others areas). We cleaned the dataset by removing non-journal items, incorrect records, and articles missing an abstract, ultimately keeping 21,486 articles for the analysis.

### 2. Selection of jargon

As for the articles, our goal was to generate a list containing the most representative biological and geological jargon used in cave-related studies. The list included both cave specific jargon (words specifically coined to describe biological or geological entities or processes restricted to caves), as well as general jargon (words used to describe general biological or geological entities, not only found in caves).

Terms were gathered from two sources. First, we screened the glossary of 10 seminal books in cave research published between 1950–2020. These included both books originally written in English as well as translations from other languages. Second, we review 21 recently published reviews discussing, among others, cave-related terminology (see Appendix S1 for the full list of these references). For each word, we manually generated all possible declinations, as well as spelling alternatives, including singular and plural forms, as well as potential adjectivations and nominalizations, regardless on whether these were found or not in the reviewed sources. For example, the noun “anchialine” was included as “anchialine”, “anchalines”, “anchihaline”, “anchihalines”, and as the general noun “anchialos”. These derivations were constructed automatically, even if this operation ended up generating some non-existing words, given that their presence would not effect the estimation of the proportion of jargon.

Through this procedure we produced an initial list of 1,333 terms, including single words (e.g., troglobiont), jargon consisting of up three words (e.g., metahaline pool, *Milieu Souterrain Superficiel*), as well as few acronyms commonly used in cave-based science (e.g., MSS).

In order to avoid flawing the analysis by confusing some jargon words with their common English homonyms (i.e., words that are spelled the same and sound the same, but have different meanings), each word in the initial list was searched in the Merrian-Webster open access English dictionary (accessed on March 2020 at https://www.merriam-webster.com/dictionary/). Words with common English homonym equivalents (e.g., “runner”) were kept in the glossary but excluded from subsequent analyses. The complete list of the sources, as well as the list of terms, including their category, inclusion of the analyses, and general definitions is in the Appendix S1.

### 3. Statistical analyses

We conducted all analyses in R (R Core Team 2018).

#### *Proportion of jargon in the title and abstract*

We focused the analyses on the title and abstract of each article, because these are the two elements that showcase it to a broader audience (Fox and Burns 2015, Plavén-Sigray et al. 2017, Bowman and Kinnan

2018, Murphy et al. 2019). Furthermore, abstracts are assumed to reflect the overall writing style of entire articles (Vinkers et al. 2015, Plavén-Sigray et al. 2017, Mammola 2020). We expressed jargon use as the proportion between the number of unique jargon terms and the total number of unique words used in the title and abstract (hereinafter 'proportion of jargon').

We processed titles and abstracts so that their total number of unique words could be counted. We first cleaned the string of text of each title and abstract using general functions for common string operations available in the R package 'stringr' (Wickham 2019), which removes punctuation, extra spaces, and transforms all letters to the lower case. In parallel, we used the function *singularize* in the R package 'SemNetCleaner' (Christensen and Yoed 2019) to make all word singular, thereby not counting plural forms when estimating the number of unique words in each abstract and title. This function automatically changes words to their singular form based on inverse of the grammar rules found in a standard blog on grammar (see Christensen and Yoed 2019 for details).

Once cleaned, we automatically counted the unique usage of jargon words in each title and abstract by matching the strings of words. To count jargon expressions made up of two or three words (e.g., "aphyletic troglophile" or "subterranean sampling device"), we also selected all possible pairs and triplets of contiguous words in the title and abstract using an R function designed *ad hoc* for this study. In calculating the proportion of jargon, we only considered unique usage of words. For example, if the word 'troglobite' appeared four times in an abstract, we counted it only once. This procedure allowed us to avoid inflating the proportion of jargon by counting the same jargon multiple times if its singular, plural, or declined forms appeared in the same title or abstract. Likewise, when a jargon words was included in a combined expression (as in "aphyletic troglobite"), we only counted it once (as "aphylotic troglobite" only and not again as "troglobite").

### Regression models

To test if an overuse of jargon in the title and abstract affects citations (Figure S1a), we conducted regression-type analyses following Zuur & Ieno's (2016) general protocol. We initially conducted data exploration (Zuur et al. 2009) checking the distribution of each variable and relationships among variables. We removed four outliers from articles citations, corresponding to four articles cited over 1,500 times in the Web of Science. We excluded two outliers in the proportion of jargon in the title and abstract, corresponding to two paper with a proportion of jargon above 0.4 for the title and above 0.08 for the abstract. Since there were almost no articles older than 28 years, we also excluded articles published before 1991 from the database.

Following Mammola et al. (2020), we used a generalized additive model to predict the expected pattern of citations over article age, and expressed the number of citations as the Pearson residuals from the curve (Figure S1b). This was a necessary step since we compared the citation of articles published in different years. Since citations are counts, we used a Poisson distribution. We constrained the generalized additive model to two knots, so to minimize tendency variations in the smoothers after it reached a plateau in the predicted number of citations (at 10 years; Figure S1b).

We used a generalized additive mixed models with a Gaussian distribution to test for the effect of jargon use on article citation residuals. The use of generalized additive mixed models allowed us to deal with non-linearities in the observed patterns and to account for dependency structures in the data. We constructed two separate models for the proportion of jargon in the title and abstract. We fitted models with the 'gamm4' R package (Wood and Scheipl 2017), using univariate penalized cubic regression spline smooths (Wood 2017). The optimum amount of smoothing was estimated trough generalized cross-validation. We included publication year (labelled as 'PY' in Web of Science) and article Research Area ('SC' in Web of Science) as random terms in a two-nested levels random-intercept structure (1 | PY/SC). This allowed us to consider data non-independence, namely the fact that articles belonging to the same subject area may use a more similar jargon than expected from random, and the variation of jargon use through time. Note that p-values for smoother terms represent approximate significance based on F statistic.

We are aware that the explanation of statistical analyses probably contains more jargon that most of the article' titles and abstracts considered in this work. Yet, we believe that only few readers interested in these methodological nuances will end up reading these technicalities in the supplementary materials.

### SUPPLEMENTARY FIGURES

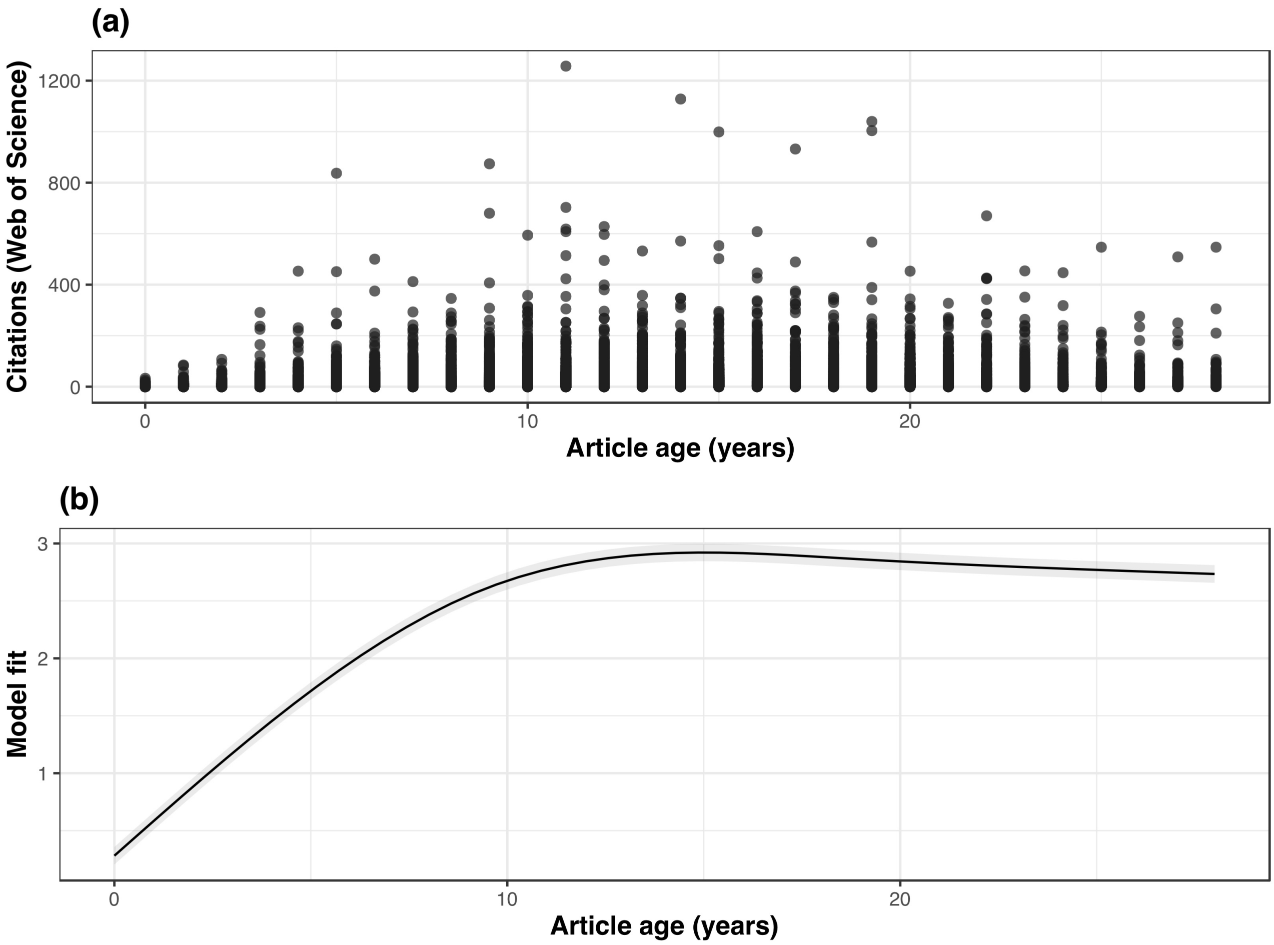


**Figure S1. Distribution of citations and their normalization by article age. a**) Distribution of citations among the articles considered in this study. Four outliers, corresponding to articles cited over 1,500 times in the Web of Science, are not shown in the graph. **b**) Expected citations over article age as predicted with a Poisson generalized additive model. To normalized the number of citations for each article by its age, we expressed citations as the Pearson's residuals from the regression curve.
