## Appendix S1 for "Specialized terminology limits the reach of new scientific knowledge"

### GLOSSARY OF THE WORDS

### 1. List of jargon

List of all the jargon terms gathered for the analyses. The references used to extract the definitions are at the bottom of the section. The definitions from the Merriam & Webster online English dictionary are included, when they were available (Searches in March 2020). Asterisks (*) indicate that the terms have been discarded from the analyses because represent homonyms to commonly used English words. The words are presented organized according to different topics within Biology and Geology, but this organization is merely indicative and it had no effect on the statistical analyses.

#### 1.1. Geological jargon

##### *1.1.1. General geological jargon*

AA lava (*derivations:* aa lava, aa lavafield)

> *Specialized definitions.* High-viscosity, lumpy lava flow [1] | Jagged, irregular volcanic rock formed from the solidification of highly viscous lavas [2,3.]

Anticline

> *Merriam & Webster:* an arch of stratified rock in which the layers bend downward in opposite directions from the crest.
>
> *Specialized definitions:* fold that is convex upward [2].

Colluvium (*derivations:* colluvia, colluvial)

> *Merriam & Webster:* rock detritus and soil accumulated at the foot of a slope.
>
> *Specialized definitions:* unconsolidated sediments deposited at the base of hill-slopes or in depressions through sheet wash or downslope creep; they may consist of silt, sand, gravel and rocks[1] | loose bodies of sediments that have been deposited of build up at the bottom of a low-grade slope, transported by gravity [3] | Colluvial: Comprised of or related to colluvium [1].

Dedolomitization (*derivations:* dedolomitisation)

> *Specialized definitions:* Replacement of Mg ions by ions of Ca in dolomites during the course of some chemical reactions that may occur during diagenesis and weathering [2].

Diastrophism

> *Merriam & Webster:* Equivalent to tectonism. The process of deformation that produces in the earth's crust its continents and ocean basins, plateaus and mountains, folds of strata, and faults.
>
> *Specialized definitions:* it designates tectonism (orogenic, epirogenetic and tafrogenetic tectonics), affecting the terrestrial crust and resulting in the formation of geological basins, mountain chains, folds, faults, fractures, joints, etc. [1].

### Appendix S1

#### Englacial

*Merriam & Webster:* embedded in a glacier.

*Specialized definitions:* In the body of the glacier [2].

#### Eogenetic

*Specialized definitions:* Means young or youthful. Eogenetic rock is rock that has not been greatly altered since its deposition; eogenetic karst is the karst developed on such rock [2].

#### Epigenetic cave

*Merriam & Webster (epigenetic):* of a deposit or structure, formed after the laying down of the enclosing rock.

*Specialized definitions:* Caves formed by meteoric water, primarily in the vadose zone [1].

#### Epikarst

*Specialized definitions:* the highly porous uppermost zone [4] | boundary region between the soil and rock in the karst, usually honeycombed with small fractures, solution pockets, and solutionally widened trenches [3] | a weathered zone of enhanced porosity near the surface, at the soil/ bedrock contact of many karst landscapes [1] | upper section of the percolation area of the karst between the unconsolidated material (soil, sediment, and vegetation debris) and the carbonate rock that is partly saturated with water [5] | the boundary region between soil and rock in karst, usually honeycombed with small fractures, solution pockets, and solutionally widened trenches (cutters or grikes) [2].

#### Euhedral

*Merriam & Webster:* (=idiomorphic) having the proper form or shape—used of minerals whose crystalline growth has not been interfered with.

*Specialized definitions:* a crystal with well-formed faces (as opposed to anhedral crystal with no faces) [2].

#### Glacioeustatic

*Specialized definitions:* The change in sea level worldwide that occurs when ice sheets grow during a glacial cycle (sea level falls) or ice sheets melt during an interglacial cycle (sea level rises) [2].

#### Kringing

*Specialized definitions:* a method for calculating a predicted value for a site in the study region that has not been sampled. The method uses a weighted mean of the nearby observed values to interpolate a value at the unobserved site. The weights are functions of the covariances between the observed sites and the unobserved site [2].

#### Lithostatic pressure

*Specialized definitions:* The pressure exerted on a rock or fluid buried deep within the Earth by overlying rocks [2].

#### Macropore

*Merriam & Webster:* a pore in soil of such size that water drains from it by gravity and is not held by capillary action.

*Specialized definitions:* a pore in soil f such size that water drains from it by gravity and is not held by capillary action. They are typically create by burrowing activities of animals and plant roots [3].

Nappe

*Merriam & Webster:* a large mass of rock thrust over other rocks | one of the two sheets that lie on opposite sides of the vertex and together make up a cone.

*Specialized definitions:* In mountainous ranges, a pile of terranes, which have been displaced by tectonic movements for kilometers on top of other terranes [2].

Nappe phréatique

*Specialized definitions:* layers of soil completely and permanently impregnated with water (Daubrée) [6].

Nunatak

*Merriam & Webster:* a hill or mountain completely surrounded by glacial ice

*Specialized definitions:* peripheral glacial refugia of mountain species [7].

Pedogenic

*Merriam & Webster:* the formation and development of soil.

*Specialized definitions:* process of soil development and evolution [3].

Sorption

*Merriam & Webster:* the process of sorbing; the state of being sorbed; to take up and hold by either adsorption or absorption.

*Specialized definitions:* The process of removing from solution and holding, either by absorption (physical assimilation) or adsorption (adhesion to the surface). Some compounds and dyes are much more susceptible to sorption losses than others [2].

Psammite

*Merriam & Webster:* a rock composed of sandy particles.

*Specialized definitions:* sand, components of the sediment with a diameter ranging between 0.02–2 mm [8].

Psephite

*Merriam & Webster:* coarse fragmental rock composed of rounded pebbles (as conglomerate).

*Specialized definitions:* gravel, components of the sediment with a diameter larger than 2 mm [8].

#### *1.1.2. Cave-related jargon*

Adit

*Merriam & Webster:* a nearly horizontal passage from the surface in a mine.

*Specialized definitions:* roughly horizontal passage introduced into a mine for the purpose of access the drainage [3].

Aeolian cave

*Merriam & Webster (for aeolian):* borne, deposited, produced, or eroded by the wind.

*Specialized definitions:* caves formed by wind erosion [5].

Aggrade

*Merriam & Webster:* to fill with detrital material.

*Specialized definitions:* to fill up with sediment [2].

Anastomic cave (*derivations:* anastomotic maze)

*Merriam & Webster:* relative to the union of parts or branches (as of streams, blood vessels, or leaf veins) so as to intercommunicate or interconnect.

*Specialized definitions:* caves with a series of tubes that sometimes intersect with each other many times, forming a complex grid with three-dimensional structure [5] | a maze of interconnected, curving, tubular passages, analogous to a braided pattern in river channels [2].

Anthodites

*Specialized definitions:* speleothem composed of clusters of needle or quill-like crystals [2].

Apron

**Merriam & Webster:* a garment usually of cloth, plastic, or leather usually tied around the waist and used to protect clothing or adorn a costume.

*Specialized definitions:* a surface that slopes down into the passage from a lava tube's wall [9].

Aven

*Specialized definitions:* a hole in a cave roof. It is the same as a shaft seen from above [5].

Baldacchino canopies

*Specialized definitions:* speleotheme consisting of an overhanging belly of calcite, which has its widest girth at the pool surface [9].

Basinal sediment

*Specialized definitions:* Sediments that accumulate in a region of low (and usually subsiding) topography [2].

Biosparite

*Specialized definitions:* a coarse-grained limestone with abundant fossils [2].

Blowhole (*derivations:* blow hole)

*Merriam & Webster:* a hole or fissure in rocks along a shore through which incoming waves force air to rush upward or water to spout intermittently.

*Specialized definitions:* A generally round hole in the ground ranging in diameter from a few tens of centimeters to one or two meters, connecting with a generally smooth-walled vertical tube of similar diameter, which may or may not descend into an accessible cave chamber. A blowhole is so named because it almost always connects with numerous small voids into which of from which atmospheric pressure changes induce air-flow (wind), sometimes of considerable strength [2].

Blue hole (*derivations:* bluehole)

*Specialized definitions:* term widely used in the Caribbean as an equivalent to sinkhole full of water.

*Boneyard

*Merriam & Webster:* a place where worn-out or damaged objects (such as cars) are collected to await disposal.

*Specialized definitions:* a type of cave passage consisting of multiple chambers interconnected by smaller openings forming a three-dimensional, swiss-cheese-like maze [2].

Braided maze

*Merriam & Webster:* made by intertwining three or more strands.

*Specialized definitions:* while many lava tubes consist primarily of a single conduit, it is not uncommon to have areas where passages branch and rejoin. Braiding occurs most actively near the leading portions of lava flows, and occurs because accretion of cooling lava occurs faster than downcutting or erosion. Hence, braiding occurs more often on the lower-gradient areas of the tube, that is, where the surface that the lava is flowing over is less steep [9].

Branchwork cave

*Specialized definitions:* A cave formed by underground streams that converge in a branching pattern, with tributaries joining downstream to form progressively fewer but (usually) larger passages [2].

Caliche

*Merriam & Webster:* a crust of calcium carbonate that forms on the stony soil of arid regions.

*Specialized definitions:* hardened deposit of calcium carbonate. Caliche occurs worldwide, generally in arid of semiarid regions [3].

Caprock

*Specialized definitions:* an impermeable rock layer, such as sandstone, covering soluble carbonates and restricting erosion [4] | an insoluble rock unit overlying soluble, cave-bearing strata. A typical caprock consists of sandstone or shale [2].

Casimba

*Specialized definitions:* small sinkholes in Cuba.

Cave balloon (*derivations:* *balloon)

*Specialized definitions:* virtual cave: small, gas-filled pouch usually made of hydromagnesite [9].

Cave pearls (*derivations:* *pearls)

*Merriam & Webster:* a small smooth round concretion of carbonate of lime found in limestone caves.

*Specialized definitions:* concentric concretion found in shallow cave pools. They can be spherical, as in these photos, or cylindrical, elliptical, and even cubical [9].

Cave popcorn (*derivations:* *popcorn)

*Specialized definitions:* popcorn is a common name for a very common type of coralloid. It can be recognized by its gregarious nature and knob-like shape, the latter of which it owes to its concentric layering of microcrystalline calcite [9].

Cave runner (*derivations:* *runner)

*Specialized definitions:* forms of tubular lava stalactite formed along a surface instead of free-hanging [9].

Cave shield (*derivations:* *shield)

*Specialized definitions:* structures formed from calcite-rich seep water under hydrostatic pressure is forced from tiny cracks in a cave wall, ceiling or occasionally, floor [9].

Cenote

*Merriam & Webster:* a deep sinkhole in limestone with a pool at the bottom that is found especially in Yucatán.

*Specialized definitions:* flooded, natural depression carved out in limestone with a collapsed ceiling [5] | Refers to places where bedrock solution has led to the collapse of surface limestone, giving access to the water table [2].

*Clinker

*Merriam & Webster:* a brick that has been burned too much in the kiln | stony matter fused together.

*Specialized definitions:* the surface of a aa lava is covered with layer of partly loose, very irregular fragments, commonly known as clinker, formed as the lava cools [3].

Conulites

*Specialized definitions:* conulites are the "splash cups" that form on certain cave floors beneath energetic ceiling drip sites [9].

Coralloid

*Merriam & Webster:* relative to corals.

*Specialized definitions:* term use to refer to all manner of knobby, globular, button-like, coral-like, or botryoidal type formations that can form either above or below water [9].

Cupola

*Merriam & Webster:* a rounded vault resting on a usually circular base and forming a roof or a ceiling | a small structure built on top of a roof.

*Specialized definitions:* dome-like heightening in a lava tube's ceiling, sometimes found in lower-level passages beneath windows that have sealed off, or caused by breakdown from a spot in the ceiling [9].

Cutbank

*Merriam & Webster:* a steep bare slope formed typically by stream erosion.

*Specialized definitions:* a concave section of a tube wall formed on the outside edge of a meandering passage, eroded on the outside side edge of a meandering passage, where the erosive force is greatest because velocity and turbulence are higher here [9].

Donga

*Merriam & Webster:* (chiefly in Africa) a narrow steep-sided ravine formed by water erosion but usually dry except in the rainy season.

*Specialized definitions:* shallow, generally circular, closed depression several meters deep and hundreds of meters across, with a flat clay-loam floor and very gently sloping sides [2].

*Drapery

*Merriam & Webster:* a decorative piece of material usually hung in loose folds and arranged in a graceful design | hangings of heavy fabric for use as a curtain.

*Specialized definitions:* cave formations, draperies are deposited from calcite-rich solutions flowing along an overhung surface. Surface tension allows these solutions to cling to a wall or sloping ceiling as they stream slowly downward [9].

Dripstone

*Specialized definitions:* Any formation resulting from the deposition of calcite, generally by percolating water [2].

Estavelle

*Merriam & Webster:* calcium carbonate in the form of stalactites or stalagmites.

*Specialized definitions:* a conduit opening to the surface that functions as a spring in high-flow conditions and as a stream sink in low-flow conditions [2].

Flank-margin cave

*Specialized definitions:* cave formed in young sea coast limestone in the freshwater-seawater mixing zone, along the flanks of the freshwater lens [2,3].

Flashine

*Specialized definitions:* the rapidity with which a water table position (and related flow and chemical conditions) changes within a karst aquifer in response to an input of water from a storm [2].

Flowstone

*Merriam & Webster:* calcite deposited by a thin sheet of flowing water usually along the walls or floor of a cave.

*Specialized definitions:* rocks composed of sheetlike deposits of calcite or other carbonate minerals, formed where water flows down the walls or along the floors of a cave [9].

*Folia

*Merriam & Webster:* one of the lamellae of the cerebellar cortex.

*Specialized definitions:* cave formation found near or just below a water line, which may not be evident as the water table may have long since dropped below the level of the cave. Found on ceiling and walls, they resemble inverted rimstone dams. They are thought to form primarily near the top of the water table, and are associated with a declining water level. Like shelfstone, they probably form from precipitates on water surfaces that accrete to walls. As the water lowers, more calcite is deposited underneath, forming a flat surface [9].

Glaze

*Merriam & Webster:* a smooth glossy or lustrous surface or finish.

*Specialized definitions:* smooth, thin, metallic-looking coating over the often darker, coarser basalt forming the walls of lava tubes brainstorm [9].

Gour

*Specialized definitions:* a type of cave formation in the form of a stone dam, also known as rimpool | little ponds surrounded by walls of stalagmites [6].

Grotto

*Merriam & Webster:* cave.

*Specialized definitions:* section of a cave that is well decorated with calcite or is otherwise aesthetically impressive [5].

*Gutter

*Merriam & Webster:* a trough along the eaves to catch and carry off rainwater.

*Specialized definitions:* an elongated depression running along the passage between the wall and a levee [9].

Helictites

*Merriam & Webster:* an irregular stalactite with branching convolutions or spines.

*Specialized definitions:* contorted depositional speleothems which grow in any direction, seemingly defying gravity [9].

Hypogenic (*derivations:* hypogenic caves, hypogenetic, hypogenetic speleogenesis)

*Merriam & Webster:* of, relating to, or constituting hypogene action or crystallization.

*Specialized definitions:* refers to an origin deep beneath the Earth's surface [3,4] | Hypogenetic cave: caves formed by rising groundwater or production of deep-seated solutional aggressiveness [1]| formed by acids generated partly or entirely at depths below the surface [2] | Hypogenetic speleogenesis: The formation of solution-enlarged permeability structures by waters ascending to a cave-forming zone from below, where deeper groundwaters in regional or intermediate flow systems interact with shallower and local groundwater flow systems [1].

Hypoxic cave

*Merriam & Webster:* (cave) characterized by reduced levels of dissolved oxygen.

*Specialized definitions:* cave with reduced oxygen partial pressure (0–15 kPa).

Interbedded

*Merriam & Webster:* a typically thin layer of one kind of sedimentary material located between layers of another kind.

*Specialized definitions:* tectonically deformed bedding, normally due to deformation during folding of the beds [2].

Intracratonic

*Specialized definitions:* a broad structural subsidence of part of the Earth's continental crust. Such basins are the sites of considerable sediment accumulation [2].

Karren

*Merriam & Webster:* a ribbed and fluted rock surface resulting at least in part from differential solution.

*Specialized definitions:* German word that means fissures or furrows [5] | Weathered and exposed limestone surfaces found in karst regions and consisting of different forms of rock pinnacles separated by deep grooves [1] | Minor forms of karst, due to solutional sculpturing of rock surfaces of grunderground [2].

Karst-siphon

*Merriam & Webster:* (siphon) tube bent to form two legs of unequal length by which a liquid can be transferred to a lower level over an intermediate elevation by the pressure of the atmosphere in forcing the liquid up the shorter branch of the tube immersed in it while the excess of weight of the liquid in the longer branch when once filled causes a continuous flow.

*Specialized definitions:* a completely flooded section within a karstic system

Keyhole passage

*Specialized definitions:* when a phreatic (elliptic) conduit has been incised by a free surface stream cutting a canyon, the cross-section of the passage is very similar to a large. keyhole [2].

Lapies (*derivations*: lapiaz)

*Merriam & Webster:* grooves and ridges formed on a rock surface by solution of limestone.

*Specialized definitions:* weathered and exposed limestone surfaces found in karst regions and consisting of different forms of rock pinnacles separated by deep grooves [1].

Laterite

*Merriam & Webster:* a residual product of rock decay that is red in color and has a high content in the oxides of iron and hydroxide of aluminum.

*Specialized definitions:* soil types rich in iron and aluminum, formed in wet tropical areas. Nearly all laterites are rusy-red because of iron oxides [3].

Lava blades

*Specialized definitions:* projections from the wall that tend to be parallel and regularly spaced [9].

Lavaball

*Specialized definitions:* cave formations are formed from pieces of breakdown that float and roll along in a stream of moving lava, accreting mass as they move [9].

Levee

*Merriam & Webster:* a continuous dike or ridge (as of earth) for confining the irrigation areas of land to be flooded.

*Specialized definitions:* as a stream of lava moves through an existing tube, the sides tend to cool first and create a free-standing, vertical remnant along the edge. Its height is increased by splashing and frothing, and the top is often irregular [9].

Looping passage

*Specialized definitions:* phreatic conduit going up and down. In many cases, several generations of conduits cross each other and form a network of “looping” conduits [2].

Mammillaries

*Specialized definitions:* carbonate coatings that form underwater in cave pools whose water is super-saturated with calcium carbonate [9].

Marginal cave

*Specialized definitions:* see flank-margin cave.

Maze cave

*Specialized definitions:* a cave composed of a complex grid of intersecting passages, usually with many closed loops [2].

Moonmilk

*Specialized definitions:* a white, plastic calcareous cave deposit composed of calcite, huntite, or magnesite [3].

Oosparite

*Specialized definitions:* a coarse-grained limestone with accretionary particles (ooliths) [2].

#### Pahoehoe

*Merriam & Webster:* cooled hard lava marked by a smooth often billowy shiny surface —contrasted with aa.

*Specialized definitions:* volcanic rock with smooth ropy surface formed from the solidification of fluid lavas [2–4] | low-viscosity lava that flows like a river ("rope lava") [1].

#### Paleodrainage

*Specialized definitions:* a drainage basin that has subsequently been altered [4] | the no longer active network of definable channels formerly occupied by a river or stream which, under less arid conditions, constituted the active surface catchment of the region [2].

#### Paleokarst

*Specialized definitions:* any geological evidence of former karstic activity [5].

#### Parafluvial

*Specialized definitions:* area of the bankfull channel that is to some extend annually scoured by flooding, and is thus lateral to the normal stream channel [3].

#### Phreatic loop

*Merriam & Webster:* of, relating to, or being groundwater.

*Specialized definitions:* phreatic conduit going up and down. In many cases, several generations of conduits cross each other and form a network of "looping" conduits [2].

#### Phreatic passage

*Merriam & Webster:* of, relating to, or being groundwater.

*Specialized definitions:* cave passages formed below the water table [2].

#### Phytokarst

*Specialized definitions:* a phenomenon where speleothems or speleogens orient towards sunlight coming from a cave entrance [9].

#### Plunge pool

*Merriam & Webster:* a small deep plunge basin.

*Specialized definitions:* when lava is ponded, as in a lava lake, the top portion cools first and forms a crust. The molten material below contracts during crystallization and cooling, and the crust may collapse. In this photo a relatively round lava lake's crust has collapsed into large wedges [9].

#### Polje

*Merriam & Webster:* an extensive depression having a flat floor and steep walls but no outflowing surface stream and found in a region having karst topography (as in parts of Yugoslavia).

*Specialized definitions:* a large spring-fed karst depression with a flat floor commonly covered by river sediment [4].

#### Ponor

*Merriam & Webster*: a steep-sided sinkhole.

*Specialized definitions:* a cave formed by underground streams that converge in a branching pattern, with tributaries joining downstream to form progressively fewer but (usually) larger passages [2].

Pozzo

*Specialized definitions:* cave pit.

Project cave

*Specialized definitions:* a broad structural subsidence of part of the Earth's continental crust. Such basins are the sites of considerable sediment accumulation [2].

*Raft

*Merriam & Webster:* a floating cohesive mass.

*Specialized definitions:* a cave rafts are delicate, doily-like sheets of calcite or aragonite that occasionally grace the surface of still, supersaturated cave pools [9].

Raft cone (*derivations:* cave rafts)

*Specialized definitions:* raft cones are heaps of sunken "cave rafts" [9].

Ramiform maze

*Specialized definitions:* a maze cave consisting of interconnected rooms and spongework, with passages extending outward as present or former outlet routes [2].

Regolith

*Merriam & Webster:* unconsolidated residual or transported material that overlies the solid rock on the earth, moon, or a planet.

*Specialized definitions:* unconsolidated solid material covering bedrock [3].

Rimstone

*Merriam & Webster:* a calcareous deposit formed as a ring around an overflowing basin (as of a mineral hot spring).

*Specialized definitions:* a secondary deposition of calcium carbonate in the form of a basin or dam that often holds percolating water [2].

Rootsicles

*Specialized definitions:* a generic term for all forms of calcite-coated roots, whether forming a column or a stalactite [9].

Saltpeter

*Merriam & Webster:* potassium nitrate.

*Specialized definitions:* white crystalline substance, usually composed of potassium nitrate, that was used during the nineteenth century to produce gunpowder and for medicinal purposes [5].

*Scallop

*Merriam & Webster:* any of numerous marine bivalve lamellibranch mollusks (family Pectinidae) that have a radially ribbed shell with the edge undulated and that swim by opening and closing the valves.

*Specialized definitions:* asymmetrical, cuspate oystershell shaped dissolution depressions in cave walls [2].

Scallop dominant discharge

*Specialized definitions:* the part of the hydrological regime represented by a scallop population [2].

Shelfstone

*Specialized definitions:* ledge or projection extending from the edge of a cave pool or attached to a speleothem dipped in a cave pool [9].

Show cave (*derivations:* show-cave, showcave)

*Specialized definitions:* caves which have man-made improvements that allow a person to tour the cave without caving equipment. The cave is managed and fees are often charged [2].

Showerhead

*Merriam & Webster:* a fixture for directing the spray of water in a bathroom shower.

*Specialized definitions:* rare type of stalactite generally found in tropical caves, which they sprout from the ceiling at a seep site [9].

Slipbank

*Specialized definitions:* apron, or a surface that slopes down into the passage from a lava tube's wall [9].

Soda straw

*Specialized definitions:* hollow, elongate, generally translucent tubes of calcite equal in diameter to the water drops conducted along their length [9].

Spar

*Merriam & Webster:* a stout pole.

*Specialized definitions:* a general term used to refer to crystals where the crystal faces are readily discernible. In caves, spar is a depositional deposit, usually made of calcite or gypsum, but sometimes of less common (to caves) minerals such as barite, fluorite, halite, or quart [9].

Spathies

*Specialized definitions:* soda straws that are formed from aragonite, rather than calcite, and differences in the crystal structure of these two minerals creates a different morphology [9].

Spatial subsidy

*Specialized definitions:* when resources from one system (e.g., surface) are transferred to another adjoining system (e.g., caves) [3,4].

Spelaean (*derivations*: spelean)

*Merriam & Webster:* dwelling or occurring in a cave.

*Specialized definitions:* pertaining to caves [5].

Speleogen

*Specialized definitions:* part of the bedrock the cave is formed in, that has been sculpted by erosion or dissolved into distinct interesting shapes [9].

Speleothanic zone

*Specialized definitions:* the upper most zone of karst, one of rapid destruction, according to Šušteršič [3].

Speleothem

*Merriam & Webster:* a cave deposit or formation.

*Specialized definitions:* a mineral deposit in a cave; popularly known as formations [2,4] | structure that is generated in a cave by the deposition of minerals from water [5].

Speleogen

*Specialized definitions:* eroded bedrock features in caves such as scallops, ceiling and floor channels, and bedrock projections and protrusions [2].

Splattermite

*Specialized definitions:* informal name that cavers use to describe a peculiar type of stalagmite featuring platy, upright protrusions [9].

Spongework maze

*Merriam & Webster:* an irregular pattern of very small interconnecting cavities sometimes produced by solution in cave walls.

*Specialized definitions:* a maze cave consisting of interconnected voids like those in a sponge, usually formed by the solutional enlargement of intergranular pores [2].

Stalagmite

*Merriam & Webster:* a deposit of calcium carbonate like an inverted stalactite formed on the floor of a cave by the drip of calcareous water.

*Specialized definitions:* upward-growing, massive calcite mounds deposited from drip water [9].

Stegamite

*Specialized definitions:* formations consisting of black calcite that appear as "ridges" along a cave floor. It appears that they are formed by water forced through cracks or joints [9].

Streamsink

*Specialized definitions:* a place where a surface stream enters a cave [1].

Suffosional cave

*Specialized definitions:* caves produced by sediments flushed by stormwaters [5].

Sumps

*Merriam & Webster:* a pit or reservoir serving as a drain or receptacle for liquids,

*Specialized definitionsa* flooded section of a cave [5] | with respect to caves, a sump is a place where the ceiling of a cave passage descends below water level [2].

Swallet

*Merriam & Webster:* an underground stream.

*Specialized definitions:* hole into which a stream flows [4].

*Syphon

*Merriam & Webster:* tube bent to form two legs of unequal length by which a liquid can be transferred to a lower level over an intermediate elevation by the pressure of the atmosphere in forcing the liquid up the shorter branch of the tube immersed in it while the excess of weight of the liquid in the longer branch when once filled causes a continuous flow.

*Specialized definitions:* section of a cave passage that is completely filled with water [1].

Tafoni

*Specialized definitions:* roughly hemispherical hollows weathered in rock either at the surface or in caves [2].

Talus cave

*Merriam & Webster:* (cave formed in) a slope formed especially by an accumulation of rock debris.

*Specialized definitions:* openings between piles of boulders that are sufficiently large to allow a human being to pass through them [5].

Tectonic cave

*Merriam & Webster:* of or relating to tectonics.

*Specialized definitions:* cave formed by ground movement, mostly landsliding in jointed rocks. Tectonic caves do not depend on dissolution for their formation [5].

*Trays

*Merriam & Webster:* an open receptacle with a flat bottom and a low rim for holding, carrying, or exhibiting articles.

*Specialized definitions:* unusual cave formation formed from clusters of popcorn which ends in a flat-bottomed surface [9].

Vadose (*derivations:* vadose cave, vadose water, vadose zone)

*Merriam & Webster:* of, relating to, or being water or solutions in the earth's crust above the permanent groundwater level.

*Specialized definitions:* zone above the water table in which water moves by gravity and capillarity. Water does not fill all the opening and does not build up pressure greater than the atmospheric [3,4] | (vadose cave): a vadose (or unsaturated) cave is one that underwent most of its development above the water table [9] | (vadose water): water migrating vertically through the unsaturated zone above the water table [2] | (vadose zone): the zone above the water table in which water moves by gravity and capillarity. Water does not fill all the openings and does not build up pressures greater than atmospheric [2,4] | the unsaturated zone of the groundwater environment between the surface and the phreatic—saturated zone [1].

Vrujla

*Specialized definitions:* in Croatia, term applied to submarine springs [10].

Vuggy

*Merriam & Webster:* a small unfilled cavity in a lode or in rock.

*Specialized definitions:* refers to rocks containing cavities, voids or large pores [1].

Zone of aeration

*Specialized definitions:* equivalent to vadose zone (see above)

#### *1.1.3. Aquifer-related jargon*

Aquiclude

*Merriam & Webster:* a geologic formation or stratum that confines water in an adjacent aquifer.

*Specialized definitions:* body of relatively impermeable rock or sediment acting as a boundary to an aquifer [3]

Aquitard

*Merriam & Webster:* a geologic formation or stratum that lies adjacent to an aquifer and that allows only a small amount of liquid to pass.

*Specialized definitions:* geological formation of a rather impervious and semi-confining nature that transmits water in a very slow rate compared to an aquifer [2,3]

Artesian

*Merriam & Webster:* (artesian) involving, relating to, or supplied by the upward movement of water under hydrostatic pressure in rocks or unconsolidated material beneath the earth's surface.

*Specialized definitions*: originally describing wells made in the French province of Artois in the eighteenth century, such that a perpendicular boring into a synclinal fold or basin of the strata produced a constant supply of water rising spontaneously to the surface of the ground. Thus, it is applied to water obtainable by artesian boring [11].

Artesian well

*Merriam & Webster:* a well in which water is under pressure.

*Specialized definitions:* wells made in the French province of Artois in the eighteenth century, such that a perpendicular boring into a synclinal fold or basin of the strata produced a constant supply of water rising spontaneously to the surface of the ground. Thus, it is applied to water obtainable by artesian boring [5]

Calcrete

*Merriam & Webster:* a limestone formed by the cementation of soil, sand, gravel, shells, by calcium carbonate deposited by evaporation, or by the escape of carbon dioxide from vadose water.

*Specialized definitions:* carbonate deposits that form in the vicinity of the water table as a result of evaporation [3] | a deposit, often nodular, of calcium carbonate formed in the soil[2].

Confined aquifer

*Specialized definitions:* an aquifer where groundwater is under pressure in a bed or stratum confined by less permeable rocks or sediment above it. A head in such an aquifer lies above the bottom of the confining bed [2].

Connate

*Merriam & Webster:* entrapped in sediments at the time of their deposition.

*Specialized definitions:* water trapped in sedimentary rock during its deposition [4].

Epiphreatic

*Specialized definitions:* lowest level of unsaturated zone, immediately above ground (phreatic) water [4].

Euhaline (*derivations*: euhaline pool)

*Specialized definitions:* marine water [2] | (euhaline pools): ponds containing marine water

Ghyben-herberg lens

*Specialized definitions:* freshwater lens on top of saltwater; its depth below sea level is approximately 40 times the height of the water table above sea level [4].

Halocline

*Merriam & Webster:* a usually vertical gradient in salinity (as of the ocean)

*Specialized definitions:* a zone, present in anchialine caves, in which there are rapid vertical changes in the salinity [2].

Hyperhaline (*derivations*: hyperhaline pool)

*Specialized definitions:* waters with salinity higher than 60–80 ppm [12] | (hyperhaline pool): pools with waters with salinity 60–80 ppmm [13].

Mesohaline

*Specialized definitions:* waters with a salinity ranging between 5–18 ppm [12].

Metahaline

*Specialized definitions:* waters with a salinity ranging between 40 and 60–80 ppm [12].

Metahaline anchialine pool

*Specialized definitions:* pools with salinities up to about twice seawater [13] | Pools with salinities elevated above the euhaline values [2].

Meteoric water

*Merriam & Webster:* (water) of, relating to, or derived from the earth's atmosphere

*Specialized definitions:* water from the earth surface water cycle, in contrast to endogenic or hypogenic water that originates from the mantle [1] | meteoric means relating to the weather, so meteoric water is water derived from rain. Meteoric water may then percolate into the ground. Meteoric groundwater is thus distinguished from other types of groundwater, such as those of thermal origin or volcanic origin [2].

Mixing zone

*Specialized definitions:* the boundary between fresh and sea water in an aquifer when there is a gradual change from one to the other [2].

Mixoeuhaline

*Specialized definitions:* waters with a salinity ranging between 30–40 ppm [12].

Mixohaline

*Specialized definitions:* brackish water [2].

Mixomesohaline

*Specialized definitions:* somewhat equivalent to mesohaline, waters with a salinity ranging between 5–18 ppm [8].

Mixooligohaline

*Specialized definitions:* somewhat equivalent to oligohaline waters with a salinity ranging between 0.5–5 ppm [8].

Mixopolyhaline

*Specialized definitions:* somewhat equivalent to polyhaline, waters with a salinity ranging between 18–30 ppm [8].

Oligohaline

*Specialized definitions:* waters with a salinity ranging between 0.5–5 ppm [12].

Perched aquifer

*Specialized definitions:* aquifer isolated from the surface by a layer of impermeable rocks

Phreatic zone

*Merriam & Webster:* of, relating to, or being groundwater

*Specialized definitions:* below the groundwater table; below the unsaturated zone [2–4] | phreatic Refers to an underground natural source of water. The phreatic zone is also the zone of saturated rock below the water table [5] | the zone below the water table where all spaces are saturated with water [1].

Polyhaline (*derivations:* polyhaline pool)

*Specialized definitions:* waters with a salinity ranging between 18–30 ppm [12].

Porous media aquifer

*Specialized definitions:* non-karst aquifers in which water generally moves slowly through small spaces between grains, as in sandstone, for example, in contrast to the rapid conduit flow common to well-developed karst aquifer [2].

Potentiometric surface

*Specialized definitions:* surface representing the level to which underground water confined in pores and conduits would rise if intersected by a borehole [2].

Pycnocline

*Specialized definitions:* interface between liquids with two different densities.

Saturated boundary layer

*Specialized definitions:* See turbulent [2].

Seepage pond

*Merriam & Webster:* (seep) a spot where a fluid (such as water, oil, or gas) contained in the ground oozes slowly to the surface and often forms a pool.

*Specialized definitions:* ponds feed by seepage springs (see seepage spring).

Seepage spring

*Merriam & Webster:* (seep) a spot where a fluid (such as water, oil, or gas) contained in the ground oozes slowly to the surface and often forms a pool.

*Specialized definitions:* small spring where water oozes out of the ground. Often associated with hypotelminorheic habitats [3,4]

Vernal pond

*Merriam & Webster:* (for vernal) of, relating to, or occurring in the spring

*Specialized definitions:* temporary pools of water that provide habitat for distinct plants and animals. They are considered to be a distinctive type of wetland usually devoid of fish, and thus allow the safe development of natal amphibian and insects species unable to withstand predation from fish [3].

### 1.2. Biological jargon

#### *1.2.1. General biological jargon*

Anophthalmy

*Specialized definitions:* Without eyes [2–4].

Chemolithoautotrophy

*Specialized definitions:* a metabolic mode of inorganic substrates which gives energy to an organism from light-independent chemical reactions and enables biochemical fixation of carbon dioxide for most or all organism's carbon requirements [1].

Chemolithotrophy

*Specialized definitions:* process of extracting nutrients from the substrate carried out by some microbial organisms [5] | metabolic energy production from electrons obtained from the oxidation of inorganic compounds derived from rocks [1].

Cryptozoa

*Merriam & Webster:* the animals that live a cryptobiotic life among the organic debris of a forest floor.

*Specialized definitions:* all eyeless and depigmented species fall into the cryptozoa, which occur in aphoti and in some cases dimly lit habitats [3].

Epibenthos (*derivations:* epibenthic, epibenthical)

*Merriam & Webster:* (epibenthos) the fauna and flora of the sea bottom between low-water mark and the mesobenthos down to a lower limit of about 100 fathoms.

*Specialized definitions:* (epibenthic) an organism that lives on the surface of sediments at the bottom of water [1].

Epigean (*derivations:* epigeal)

*Merriam & Webster:* (*derivations:* epigeal) living on or near the surface of the ground; relating to or being the environment near the surface of the ground

*Specialized definitions:* surface environment; also applies to organisms living there [5] | an organism living above the ground, on the surface—in contrast to subterranean or below the ground organisms [1]| living on or near the surface of the ground, as opposed to the subsurface [3].

Epilithic (*derivations:* epilithical)

*Merriam & Webster:* growing upon stone or stonelike material.

*Specialized definitions:* growing on rock [4].

Epizoic (*derivations:* epizoite)

*Merriam & Webster:* living upon the body of an animal.

*Specialized definitions:* living on the surface of (another) animal without harming it [2].

Euryhaline

*Specialized definitions:* pertains to organisms that are tolerant of a wide range of salinity [2].

Exaptation (*derivations:* exaptative)

*Merriam & Webster:* a trait, feature, or structure of an organism or taxonomic group that takes on a function when none previously existed or that differs from its original function which had been derived by evolution | the condition or circumstance of possessing one or more such traits, features, or structure.

*Specialized definitions:* adaptation for one function serving for another function [3] [4] | an apparent morphological or physiological adaptation of an organism to a habitat with which it has not (yet) been in contact; also called also "pre-adaptation" [2].

Humicolous (*derivations*: humicoles)

*Specialized definitions:* animal living in humus [2,6].

Hydrogenotrophic

*Specialized definitions:* bacteria and Archaea that utilize hydrogen as an electron donor in chemoautotrophy [4].

Hyperbenthos (*derivations*: hyperbenthic)

*Specialized definitions:* within an aquatic environment, volume of water near the bottom.

Iteroparity

*Specialized definitions:* repeated reproduction within the organisms life [2].

Lapidicole (*derivations:* lapidicolous)

*Merriam & Webster:* (lapidicolous) living under a stone —used especially of an insect

*Specialized definitions:* living under a stone [3].

Lavicole

*Specialized definitions:* an organism living exclusively on barren lavas and feeding on fallout of aerial plankton [1].

Living fossil

*Merriam & Webster:* an organism (such as a horseshoe crab or a ginkgo tree) that has remained essentially unchanged from earlier geologic times and whose close relatives are usually extinct.

*Specialized definitions:*

Madicole

*Specialized definitions:* fauna and flora living on a rocky wall with a small layer of running water.

Microphthalmia (*derivations:* microphthalmous)

*Merriam & Webster:* abnormal smallness of the eye usually occurring as a congenital anomaly.

*Specialized definitions:* organism with reduced eyes relative to a surface relative [14].

Mixotroph (*derivations:* mixotrophic)

*Merriam & Webster:* (mixotrophic) deriving nourishment from both autotrophic and heterotrophic mechanisms —used especially of symbionts and partial parasites.

*Specialized definitions:* organisms that combine two kinds of energy acquisition, the autotroph and the heterotroph [2].

Muscicole (*derivations:* muscicolous)

*Merriam & Webster:* (muscicolous) growing on decaying mosses or hepatics.

*Specialized definitions:* inhabitants of mosses or hepatics [6.]

Myrmecophile (*derivations:* myrmecophilous)

*Merriam & Webster:* an organism that habitually shares an ant nest.

*Specialized definitions:* organisms that live associated to ants or their nests [6].

Neoteny (*derivations:* neotenia)

*Merriam & Webster:* retention of some larval or immature characters in adulthood | attainment of sexual maturity during the larval stage.

*Specialized definitions:* retardation of somatic development, so that sexual maturity is attained in an organism retaining juvenile characters [4] | retention of some larval or immature characters by adults in a species [1] | The presence of juvenile features in an adult animal [2].

Neuston

*Merriam & Webster:* minute organisms that float in the surface film of water.

*Specialized definitions:* collection of organisms which float on the surface of the water [6].

Normoxia

*Specialized definitions:* the condition of having a normal concentration of oxygen.

Orthogenesis

*Merriam & Webster:* variation of organisms in successive generations that in some especially former evolutionary theories takes place in some predestined direction resulting in progressive evolutionary trends independent of external factors.

*Specialized definitions:* evolution toward a 'perfect form', determined by factors internal to the organism [4] | the development of a fixed evolutionary program independently of the particular environmental or ecological conditions experienced by the species. This fix program can be similar in different lineages, resulting in parallel or convergent evolution [1] | theory of evolution toward a preordained form determined by inherent features of the initial ancestral organism [2].

Paedogenetic (*derivations:* paedogenesis)

*Merriam & Webster:* reproduction by young or larval animals.

*Specialized definitions:* reproduction by paedomorphic animals (see paedomorphism).

Paedomorphism (*derivations:* paedomorphic, paedomorphosis)

*Merriam & Webster:* of, relating to, involving, or exhibiting paedomorphosis or paedomorphism; retention in the adult of infantile or juvenile characters

*Specialized definitions:* precocious sexual maturity in an organism that is still at a morphologically juvenile stage [4].

Photopathy

*Merriam & Webster:* pronounced and usually negative photoaxis or phototropism.

*Specialized definitions:* organisms showing negative photoaxis or phototropism.

Phytophagous

*Merriam & Webster:* feeding on plants.

*Specialized definitions:* feeding on plants [4].

Progenesis

*Merriam & Webster:* precocious sexual reproduction in a trematode worm in which metacercariae or sometimes cercariae may lay eggs capable of repeating the life cycle.

*Specialized definitions:* acceleration of sexual maturation relative to the rest of development[2,3].

Progressive evolution

*Specialized definitions:* Opposite of regressive evolution. The exaggeration of morphological and behavioral characters that accompanies isolation in caves (e.g., elongation of antennae in insects).

Regressive evolution

*Specialized definitions:* The loss of morphological and behavioral characters that accompanies isolation in caves [4].

Relictualization

*Specialized definitions:* process of being separated from a parent population by some vicariant event [15].

Rhizophagous

*Specialized definitions:* feeding on roots [1].

Riparian

*Merriam & Webster:* relating to or living or located on the bank of a natural watercourse (such as a river) or sometimes of a lake or a tidewater.

*Specialized definitions:* pertaining to the banks of a river or stream [3] [4].

Saprophilic

*Specialized definitions:* animal obtaining organic matter from dead and decaying organisms[2].

Saprovore

*Specialized definitions:* a species feeding on dead or decaying organic matter [2].

Sapsucking

*Specialized definitions:* feeding on sap [1].

Scotophilia

*Specialized definitions:* the tendency to stay away from light exhibited by many hypogean organisms, including blind ones [5] | avoidance of bright light [2].

Semelparity (*derivations:* semelparous)

*Merriam & Webster:* (semelparous) reproducing or breeding only once in a lifetime

*Specialized definitions:* only one reproduction within the organism's life [2].

Stenohaline

*Merriam & Webster:* organism unable to withstand wide variation in salinity of surrounding water

*Specialized definitions:* animal bound to a strongly defined salinity value[2].

Straminicole

*Specialized definitions:* living in the litter [3].

Teneral

*Merriam & Webster:* of, relating to, or constituting a state of the imago of an insect immediately after molting during which it is soft and immature in coloring

*Specialized definitions:* insect recently emerged from a pupa and with a soft exoskeleton [2].

Termitophile

*Merriam & Webster:* an insect normally living in association with termites in their nests

*Specialized definitions:* organisms that live associated to termites or their nests [6]

Trophic autarchy

*Specialized definitions:* synonym of nutritional autarchy.

Zeitgeber

*Specialized definitions:* an external clock setter for circadian clocks [3].

#### 1.2.2 *General biological jargon*

Accidental species

*Specialized definitions:* An organism that is rarely found in caves, and when found there, it is not because the organism is making any real use of the habitat [5] | Surface dwelling species that get into caves by mistake and cannot survive in the underground environment [2] | Trogloxene, species only occurring sporadically in a hypogean habitat and unable to establish a subterranean population [16] | Animals without an ecological relationship with caves [17]

Aphyletic troglophile

*Specialized definitions:* subtroglophile, species inclined to perpetually or temporarily inhabit a subterranean habitat but is intimately associated with epigean habitats for some biological functions (daily e.g. feeding, seasonally, or during the life history e.g. reproduction) [16] [6].

Dudlich system

*Specialized definitions:* classification of subterranean species proposed by Dudlich

Edaphobiont

*Specialized definitions:* an organism living in deep soil [5] | an obligatory soil-dwelling species [3].

Edaphomorphic

*Specialized definitions:* morphological charactersitics associated with life in the soil, especially miniaturizarion, appendage reduction, and body thinning [3].

Edaphic (*derivations:* edaphon, eu-edaphic, euedaphic)

*Specialized definitions:* Refers collectively to organisms living in the soil [5].

Edaphophile

*Specialized definitions:* a species that completes its entire life cycle in the soil, but can also complete its entire life cycle elsewhere [3].

Edaphoxene

*Merriam & Webster:* of or relating to the soil.

*Specialized definitions:* a species that is regularly found in the soul but does not complete its life cycle in there [3].

Endogean (*derivations:* endogeous)

*Specialized definitions:* environment underneath the Earth's surface [5] | beneath the surface of the ground, usually taken to mean the soil [3] | Animal living in soil | true inhabitants of the soil [6].

Endogeomorphism

*Specialized definitions:* applied to the medium of the soil, or their inhabitatns [6].

Epigeomorphism

*Specialized definitions:* opposite of troglomorphism; of an animal having morphological (or other) adaptations typical of surface habitats (see troglomorphism).

Eucaval

*Specialized definitions:* equivalent to troglobionts [6].

Eutroglobiont

*Specialized definitions:* troglobiont, strongly bound to hypogean habitats [6,16].

Eutroglomorphism

*Specialized definitions:* the morphological, physiological and behavioral adaptations of eutroglophiles.

Eutroglophile

*Specialized definitions*: essentially epigean species able to maintain a permanent subterranean population (which may become troglobiotic) [6,16].

Eutrogloxene

*Specialized definitions:* trogloxene, species only occurring sporadically in a hypogean habitat and unable to establish a subterranean population [6,16].

Facultative troglophile

*Specialized definitions:* eutroglophile, essentially epigean species able to maintain a permanent subterranean population (which may become troglobiotic) [6,16].

Guanobia

*Specialized definitions:* equivalent to guanobite; organism that inhabits the guano in caves [6].

Guanobite

*Specialized definitions:* organism that inhabits and/or makes extensive use of guano as a source of food [5] | species that exclusively inhabit guano depoisit in caves, and whose entire biological cycle takes place in this substrate [3]| Species that, when in caves, exclusively inhabit guano deposits, and whose entire biological cycle takes place in this substrate [2].

Guano community (*derivations:* guanofauna)

*Specialized definitions: organisms* associated with guano piles [18].

Guanomorphism

*Specialized definitions:* the morphological, physiological and behavioral adaptations of guanobionts (see guanobionts and troglomorphism).

Guanophage (*derivations:* guanophagous, guanophagia)

*Specialized definitions:* organism that feeds on guano [5] | Animal that feeds directly on guano and/or on microorganisms (bacteria and fungi, for instance) that grow on it [2].

Guanophile

*Specialized definitions:* species that may live and reproduce both in guano piles and in other substrates in the cave environment [3]| Species that, when in caves, may inhabit and reproduce both in guano piles and in other substrates in the cave environment [2].

Guanoxene

*Specialized definitions:* species that, when in caves, may be found feeding and reproducing on guano deposits but depend on other substrates in the caves to complete their biological cycle [3]| Species that, when in caves, may be found feeding and/or reproducing on guano deposits but depend on other substrate(s) in the caves to complete their biological cycle [2]

Hemitroglobiont

*Specialized definitions:* eutroglophile, essentially epigean species able to maintain a permanent subterranean population (which may become troglobiotic) [6,16].

Hemitroglomorphism

*Specialized definitions:* the morphological, physiological and behavioral adaptations of hemitroglobionts (see Hemitroglobiont and troglomorphism) [6,16].

Hydrophilic

*Merriam & Webster:* of, relating to, or having a strong affinity for water.

*Specialized definitions:* organisms adapted to humid climates, showing low or no tolerance to dry conditions [2].

Hygropetric

*Specialized definitions:* A steep or vertical rocky surface, covered by a thin layer of moving water [3,4] | A steep or vertical rocky surface, covered by a thin layer of moving water; if outside caves, it is inhabited mostly by algae, mosses, and some aquatic insect larvae [2]| (hygropetric fauna) Association of animals which lives in the thin layer of water over rocks, which issues from seepages [6].

Hypogean

*Merriam & Webster:* growing or living below the surface of the ground.

*Specialized definitions:* subsurface or subterranean environment as opposed to the epigean one [3–5] | Subterranean, underground [1].

Hypogean environment

*Specialized definitions:* subsurface or subterranean environment as opposed to the epigean one [2–5] | Subterranean, underground [1].

Hypogeobiont

*Specialized definitions:* animals obligatorily and totally bound to the subterranean environment [19].

Hypogeomorphism (*derivations:* hypogebiomorphism)

*Specialized definitions:* analogous of troglomorphism.

Hypogeophile

*Specialized definitions:* animals only partially (in a more temporal than spatial manner) bound to the subterranean environment [19].

Hypogeous

*Merriam & Webster:* growing or living below the surface of the ground

*Specialized definitions:* analogous of hypogean.

Hypogeoxene

*Specialized definitions:* analogous of trogloxene.

Lampenflora

*Specialized definitions:* proliferation of phototrophic organisms near artificial light sources, especially in caves [3] | Plants growing in the vicinity of artificial light sources in caves [2].

Macrocavernicola (*derivations:* microcavernicolous)

*Specialized definitions:* inhabiting macrocavernous spaces. See macrocaverns.

Microclasiphile

*Specialized definitions:* adapted to superficial subterranean rocky habitats.

Neotroglobiont (*derivation:* Neotroglobite)

*Specialized definitions:* recently evolved troglobiont (see troglobiont).

Obligate troglophile

*Specialized definitions:* troglobiont, strongly bound to hypogean habitats [6,16].

Occasional cavernicole

*Specialized definitions:* trogloxene, species only occurring sporadically in a hypogean habitat and unable to establish a subterranean population [6,16].

*Occasional guest

*Specialized definitions:* trogloxene, species only occurring sporadically in a hypogean habitat and unable to establish a subterranean population [6,16].

Occasional trogloxene

*Specialized definitions:* trogloxene, species only occurring sporadically in a hypogean habitat and unable to establish a subterranean population [6,16].

Petrimadicolous

*Specialized definitions:* association of animals which lives in the thin layer of water over rocks, which issues from seepages (equivalent to hygropetric) (by Vaillant) [6].

Pholeophile

*Specialized definitions: animals that live associated to other animals burrows or nests* [6].

Pholeuonoid

*Specialized definitions:* One of the shapes of troglobiotic beetles with spindle-shaped trunk and elongated antennae and legs [2].

Phyletic troglophile

*Specialized definitions:* eutroglophile, essentially epigean species able to maintain a permanent subterranean population (which may become troglobiotic) [6,16].

Pseudotroglobiont (*derivations: pseudotroglobite*)

*Specialized definitions:* subtroglophile, species inclined to perpetually or temporarily inhabit a subterranean habitat but is intimately associated with epigean habitats for some biological functions (daily e.g. feeding, seasonally, or during the life history e.g. reproduction) [6,16].

Regular trogloxene

*Specialized definitions:* subtroglophile, species inclined to perpetually or temporarily inhabit a subterranean habitat but is intimately associated with epigean habitats for some biological functions (daily e.g. feeding, seasonally, or during the life history e.g. reproduction) [6,16].

Schiner-Racovitza system

*Specialized definitions:* Classification of subterranean animals on ecological grounds into troglobionts, troglophiles, and trogloxenes [3,4].

Speleobiont

*Specialized definitions:* analogous of troglobiont.

Speleophilous (*derivations:* speleophilic)

*Specialized definitions:* analogous of troglophile.

Subedaphophile

*Specialized definitions:* as subtroglophile, but in term of association to soil habitats [6].

Subguanophile

*Specialized definitions:* as subtroglophile, but in term of the association to guano [6].

Subtroglophile

*Specialized definitions:* species inclined to perpetually or temporarily inhabit a subterranean habitat but is intimately associated with epigean habitats for some biological functions (daily e.g. feeding, seasonally, or during the life history e.g. reproduction) [16].

Subtrogloxene

*Specialized definitions:* subtroglophile, species inclined to perpetually or temporarily inhabit a subterranean habitat but is intimately associated with epigean habitats for some biological functions (daily e.g. feeding, seasonally, or during the life history e.g. reproduction) [16].

Subtwilight zone (*derivations:* subliminal zone)

*Specialized definitions:* Let I be the total luminosity measurable at the entrance of the cave and i the intensity of light at a certain point of the cave, a subtwilight zone is the area of the cave where $1/2\ I \geq i \geq 1/4\ I$ [20].

Troglobie

*Specialized definitions:* equivalent to troglobiont [6].

Troglobiont

*Merriam & Webster:* an animal living in or restricted to caves especially; one occurring in the lightless waters of caves.

*Specialized definitions:* Obligate, permanent resident of terrestrial subterranean habitats [2–4]| An animal that lives only in terrestrial subterranean habitats [1]| Strongly bound to hypogean habitats [16] [6] | Species forming exclusively subterranean source populations, with sinkpopulations which may be found in surface habitats [17].

Troglobiota

*Specialized definitions:* same as troglofauna.

Troglobite

*Merriam & Webster:* troglobite, an animal living in or restricted to caves especially; one occurring in the lightless waters of caves.

*Specialized definitions:* Any of the organisms found in caves that display convergent phenotypes (morphological, physiological, and behavioral) such as loss of eyes and pigmentation [2,5]| An animal that lives only in terrestrial subterranean habitats [1] | troglobiont, strongly bound to hypogean habitats [6,16].

Troglodyte

*Merriam & Webster:* a member of any of various peoples (as in antiquity) who lived or were reputed to live chiefly in caves.

*Specialized definitions:* people or animals that venture into or live in caves. It is sometimes employed in the popular press [5].

Troglofauna

*Specialized definitions:* A general term covering all air-breathing (terrestrial) animals living underground in caves and in the subterranean matrix of the broad landscape [1].

Troglomorphic (*derivations:* troglobiomorphic, biotroglomorphic)

*Specialized definitions:* Pertaining to morphological and behavioural characters that are convergent in subterranean populations [2–4] | Organisms that show reduction or loss of phenotypic characteristics related to the hypogean environment [5].

Troglomorphism (*derivations:* troglobiomorphism, biotroglomorphism)

*Specialized definitions:* A general term covering all air-breathing (terrestrial) animals living underground in caves and in the subterranean matrix of the broad landscape [1].

Troglophile

*Specialized definitions:* Organism that can complete its life cycle in caves but may also do so outside of caves [5]| An animal that can live both on the surface and in subterranean habitats [1] | Species able to live and reproduce in subterranean habitats as well as in the epigean domain [2,3] | eutroglophile, essentially epigean species able to maintain a permanent subterranean population (which may become troglobiotic) [6,16] | A species forming source populations both in subterranean and surface environments, with individuals regularly migrating between these habitats, promoting gene flow [17].

Troglophile 1st category

*Specialized definitions:* eutroglophile, essentially epigean species able to maintain a permanent subterranean population (which may become troglobiotic) [6,16].

Troglophile 2nd category

*Specialized definitions:* troglobiont, strongly bound to hypogean habitats [6,16].

Trogloxene

*Specialized definitions:* Species appearing sporadically in subterranean habitats; called accidentals by some authors [3,4] | An animal that is only rarely entering caves, accidentally or in search for shelter on short periods of time [1]M | species only occurring sporadically in a hypogean habitat and unable to establish a subterranean population [6,16] | A species having source populations in epigean habitats, but that may form sink population in subterranean habitats [17].

Twilight zone (*derivations:* liminal zone)

*Merriam & Webster:* a world of fantasy or illusion.

*Specialized definitions:* Area of cave extending from the limit of green plants to darkness, where the light gradually attenuates [2] | Let I be the total luminosity measurable at the entrance of the cave and i the intensity of light at a certain point of the cave, the twilight zone is the area of the cave where $i \geq 1/2\ I$ [20].

Tychocaval

*Specialized definitions:* eutroglophile, essentially epigean species able to maintain a permanent subterranean population (which may become troglobiotic) [6,16].

Tychotroglobiont

*Specialized definitions:* trogloxene, species only occurring sporadically in a hypogean habitat and unable to establish a subterranean population [6,16].

Xenocaval

*Specialized definitions:* trogloxene, species only occurring sporadically in a hypogean habitat and unable to establish a subterranean population [6,16].

Subglacial refuge

*Specialized definitions:* a habitat under an ice sheet that preserves aquatic subterranean fauna during climate changing periods [1].

Thermal neutral zone

*Specialized definitions:* range of ambient temperatures in which the metabolic rate of a homeothermic animal is at its lowest level (basal metabolic rate) [2].

#### *1.2.3. Jargon specific to subterranean aquatic animals*

Amphibiont

*Specialized definitions:* aquatic species or population whose life cycle requires both surface water and groundwater habitats [4].

Amphibite

*Specialized definitions:* species that require both surface and hypogean waters in order to fulfill their life cycles [5].

Anchialine (*derivations:* anchihaline)

*Merriam & Webster:* having an underground connection to a larger tidal body (such as the sea) but no surface connection.

*Specialized definitions:* subterranean habitats, with more or less extensive connections to the sea, and showing noticeable marine as well as terrestrial influences [3,4].

Crenobiont

*Specialized definitions:* organisms normally found in springs and spring brooks, i.e. at the edge of the hypogean environment [5].

Crenostygobiont

*Specialized definitions:* stygobiontic organism normally found in springs and spring brooks.

Domaine phreatique terrestre

*Specialized definitions:* the complex of subterranean aquatic environments [25].

Endogaeolimnon

*Specialized definitions:* see stygon-.

Endogaeothalassocoen

*Specialized definitions:* the ecosystem in the saline groundwater [8].

Eulimnobenthal

*Specialized definitions:* biotope in the superficial benthic zone of lakes [8].

Eulimnobenthos

*Specialized definitions:* biocoenosis in the superficial benthic zone of lakes [8].

Eulimnocoen

*Specialized definitions:* ecosystem in the interstitial stygozone of lakes [8].

Eulimnon

*Specialized definitions:* ecosystem in the bottom of lakes [8].

Eulimnostygon

*Specialized definitions:* biocoenosis in groundwater bodies that are isolated from the coenological influences of surface-water [8].

Eulimnostygopsammal

*Specialized definitions:* mesopsammal of eulimnocoen, biotope in the sand interstitial groundwaters in the bottom of lakes [8]

Eulimnostygopsammon

*Specialized definitions:* mesopsammon of eulimnocoen, biocoenosis in the sand interstitial groundwaters in the bottom of lakes [8]

Eulimnostygopsephal

*Specialized definitions:* mesopsephal of eulimnocoen, biotope in the gravel interstitial groundwaters in the bottom of lakes [8]

Eulimnostygopsephon

*Specialized definitions:* mesopsephon of eulimnocoen, biocoenosis in the gravel interstitial groundwaters in the bottom of lakes [8]

Eupsammon

*Specialized definitions:* biocoenosis of animals truly adapted to live between the sand grains.

Eustygal

*Specialized definitions:* biotope in groundwater bodies that are isolated from the coenological influences of surface-water [8].

Eustygocoen

*Specialized definitions:* ecosystem in groundwater bodies that are isolated from the coenological influences of surface-water [8].

Eustygophile

*Specialized definitions:* same as eutroglophile, but referred to an aquatic organism.

Eustygopsammal

*Specialized definitions:* mesopsammal of eustygocoen, biotope in the sand interstitial groundwaters that are isolated from the coenological influences of surface-water [8].

Eustygopsammon

*Specialized definitions:* mesopsammon of eustygocoen, biocoenosis in the sand interstitial groundwaters that are isolated from the coenological influences of surface-water [8].

Eustygopsephal

*Specialized definitions:* mesopsephal of eustygocoen, biotope in the gravel interstitial groundwaters that are isolated from the coenological influences of surface-water [8].

Eustygopsephon

*Specialized definitions:* mesopsephon of eustygocoen, biocoenosis in the gravel interstitial groundwaters that are isolated from the coenological influences of surface-water [8].

Helocrene

*Specialized definitions:* spring originating from a marsh or bog (Springs Stewardship institute web page)

Hydrobiont

*Specialized definitions:* species inhabiting aquatic subterranean environments. See stygobiont.

Hygropsammon

*Specialized definitions:* biocoenosis in moist sands [8].

Hypocrene

*Specialized definitions:* A buried spring where flow does not reach the surface, typically due to very low discharge and high evaporation or transpiration (Springs Stewardship institute web page).

Hyporheos

*Specialized definitions:* The assemblage of organisms living in the hyporheic zone [1] | an ecotonal assemblage of epigean and hypogean organisms living within the interstitial space of superficial layers of riverbed sediments [2]

Hypothelminorheos

*Specialized definitions:* persistent wet spot, a kind of perched aquifer; fed by subsurface water in a slight depression in an area of low to moderate slope; rich in organic matter; underlain by a clay layer typically 5–50 cm beneath the surface; with a drainage area typically of less than 10,000 m2; and with a characteristic dark colour derived from decaying leaves which are usually not skeletonized [3,4]

Hypothelmon

*Specialized definitions:* fauna proper of interstitial aquatic habitats.

Interstitial community

*Specialized definitions:* animal community present in the interstices between grains of sand [6].

Interstitial ecosystem

*Specialized definitions:* aquatic ecosystem hosted between the sand grains [6].

Interstitial fauna

*Specialized definitions:* fauna living in the interstitial medium [6].

Interstitial habitat

*Specialized definitions:* habitat in the spaces between particles [4,5] | small spaces filled with water between grains of sand found in sediments below lakes and wetlands, gravel bars in rivers and sand below streams [1].

Interstitial highway

*Specialized definitions:* hypothesis that interstitial habitats, especially along rivers and streams, are a corridor for subterranean dispersal [3].

Interstitial medium

*Specialized definitions:* the aquatic environment present in the minute interstices between grains of sand [6].

Interstitial species (*derivations:* interstitial animal)

*Specialized definitions:* living in the aquatic habitat between the sand grains [6].

Krenon (*derivations:* crenon)

*Specialized definitions:* the "ecological system" associated with the outflow of springs [8].

Krenostygon (*derivations:* crenostygon)

*Specialized definitions:* ecosystems in the groundwater associated to the outflow of springs [8].

Krenostygopsammal

*Specialized definitions:* mesopsammal of krenostygon, biotope in the sandy deposits in the outflow of springs [8].

Krenostygopsammon

*Specialized definitions:* mesopsammon of krenostygon, biocoenosis in the sandy deposits in the outflow of springs [8].

Krenostygopsephal

*Specialized definitions:* mesopsephal of krenostygon, biotope in the gravelly deposits in the outflow of springs [8].

Krenostygopsephon

*Specialized definitions:* mesopsephon of krenostygon, biocoenosis in the gravelly deposits in the outflow of springs [8].

Limnobenthos

*Specialized definitions:* biocoenosis in the bottom of the lakes [8].

Limnocrene

*Specialized definitions:* spring where water comes out of the ground and creates a pond at the source, before flowing out slowly [3] | spring originating from a large, deep pool of water (Springs Stewardship institute web page).

Limnon

*Specialized definitions:* biocoenosis of the lakes [8].

Limnopsammon

*Specialized definitions:* biocoenosis in the sand of the lakes [8].

Limnostygobiont

*Specialized definitions:* stygobiont living in live in fresh groundwater [2].

Limnostygon

*Specialized definitions:* biocoenosis in the subterranean lakes [8].

Marine interstitial

*Specialized definitions:* referred to the spaces between the sand grains in haline aquatic environments [6].

Meiobenthos

*Specialized definitions:* biocenosis of microscopic animals living in the bottom of aquatic habitats.

Meiofauna

*Merriam & Webster:* the mesofauna of the benthos.

*Specialized definitions:* Assemblage of animals that pass through a 500 µm sieve but are retained by a 40 µm sieve [3–5] | Small benthic invertebrates that live in both marine and fresh water environments; organisms that can pass through a 1 mm mesh but will be retained by a 45 µm mesh [21,1].

Mesopsammal

*Specialized definitions:* biotopes in psammite (sand, sedimentary particles with a diameter ranging between 0.02–2 mm) [8].

Mesopsammon

*Specialized definitions:* biocenosis in psammite (sand, sedimentary particles with a diameter ranging between 0.02–2 mm) [8].

Mesopsephal

*Specialized definitions:* biotopes in psephite (gravel, sedimentary particles with a diameter larger than 2 mm) [8].

Mesopsephon

*Specialized definitions:* biocoenosis in psephite (gravel, sedimentary particles with a diameter larger than 2 mm) [8].

Microfauna

*Merriam & Webster:* minute animals especially, those invisible to the naked eye.

*Specialized definitions:* Animals with the body size up to 0.2 mm such as nematodes, rotifers and small arthropods, also covering protozoans (protist kingdom) [1].

Pedostygon

*Specialized definitions:* organisms adapted to drenched soil habitats.

Phreatobite (*derivations:* phreatobia, phreaticole)

*Specialized definitions:* (An organism found only in groundwater | organisms which populate the phreatic nappes [22].

Phreatophile

*Specialized definitions:* equivalent to stygophile [6].

Potamobenthal

*Specialized definitions:* biotope in the superficial benthic ecozone of rivers [8].

Potamobenthos

*Specialized definitions:* biocoenosis in the superficial benthic ecozone of rivers [8].

Potamon

*Specialized definitions:* ecosystem in the bottom of rivers [8].

Potamocoen

*Specialized definitions:* ecosystem in the superficial benthic ecozone of the rivers [8].

Potamostygocoen

*Specialized definitions:* ecosystem in the groundwater associated to rivers [8].

Potamostygal

*Specialized definitions:* biotope in groundwater associated to rivers [8].

Potamostygon

*Specialized definitions:* biocoenosis in groundwater associated to rivers [8].

Potamostygopsammal

*Specialized definitions:* mesopsammal of potamostygocoen, biotope in the sand interstitial groundwaters of the rivers [8].

Potamostygopsammon

*Specialized definitions:* mesopsammon of potamostygocoen, biocoenosis in the sand interstitial groundwaters of the rivers [8].

Potamostygopsephal

*Specialized definitions:* mesopsephal of potamostygocoen, biotope in the gravel interstitial groundwaters of the rivers [8].

Potamostygopsephon

*Specialized definitions:* mesopsephon of potamostygocoen, biocoenosis in the gravel interstitial groundwaters of the rivers [8].

Psammolittoral

*Specialized definitions:* psammal in the littoral zone of lakes or oceans [8].

Rheocrene

*Specialized definitions:* spring that flows from a defined opening into a confined channel (Bornhauser, 1913).

Rheophile

*Specialized definitions:* animals preferring living in fast moving waters [6].

Rhithrobenthos

*Specialized definitions:* biocoenosis in the superficial benthic ecozone of mountain streams [8].

Rhithrocoen

*Specialized definitions:* ecosystem in the superficial benthic ecozone of mountain streams [8].

Rhithron

*Specialized definitions:* the ecosystem associated with the bottom of mountain streams [8].

Rhithrostygocoen

*Specialized definitions*: ecosystem of the groundwater in the mountain streams [8].

Rhithrostygon

*Specialized definitions:* biocoenosis of the groundwater in the mountain streams [8].

Rhithrostygopsammal

*Specialized definitions:* biotope of the sand interstitial groundwater in the mountain streams [8].

Rhithrostygopsammon

*Specialized definitions:* biocoenosis of the sand interstitial groundwater in the mountain streams [8].

Rhithrostygopsephal

*Specialized definitions:* biotope of the gravel interstitial groundwater in the mountain streams [8].

Rhithrostygopsephon

*Specialized definitions:* biocoenosis of the gravel interstitial groundwater in the mountain streams [8].

Stygobiont

*Specialized definitions:* obligate, permanent resident of aquatic subterranean habitats [3,4] | An animal that lives only in groundwater [1] | Animal species inhabiting subterranean waters and occurring only in subterranean waters, either in interstitial habitats or in free water of caves or other karstic habitats [2].

Stygobiota

*Specialized definitions:* aquatic troglobiota (see above).

Stygobite

*Specialized definitions:* any hypogean organism that shows some sort of specialization to the underground aquatic environment [5] | An animal that lives only in groundwater [1] | Animal species inhabiting subterranean waters and occurring only in subterranean waters, either in interstitial habitats or in free water of caves or other karstic habitats [2].

Stygocoen

*Specialized definitions:* the ecosystem “groundwater” [23,24].

Stygofauna

*Specialized definitions: f*auna inhabiting the various types of groundwater [3,4] | Collective term for the animals inhabiting the underground water environment [5] | A general term covering all animals living in groundwater associated with caves, streams and the broad landscape [1].

Stygomorphic

*Specialized definitions:* describes an organism that displays the convergent phenotypic (morphological, physiological, and behavioral) characteristics of stygobites [5].

Stygon (*derivations:* stygios)

*Specialized definitions:* the ecosystem associated to groundwater [8].

Stygophile

*Specialized definitions:* an aquatic organism that can complete its life cycle in caves but may also do so outside of caves [5] | An animal that lives both on the surface and in groundwater habitats [1] | species that completes its entire life cycle in subterranean waters but can also complete its entire life cycle in surface waters [2,3].

Stygopsammal

*Specialized definitions:* biotope in the sand interstitial groundwaters [8].

Stygopsammon

*Specialized definitions:* biocoenosis in the sand interstitial groundwaters [8].

Stygopsephal

*Specialized definitions:* biotope in the gravel interstitial groundwaters [8].

Stygopsephon

*Specialized definitions:* biocoenosis in the gravel interstitial groundwaters [8].

Stygos

*Specialized definitions:* generic name for subterranean water [23].

Stygoxene

*Specialized definitions:* An organism that can be found accidentally in the hypogean environment [5]| An animal that is only rarely living in groundwater habitats, accidentally or in search for shelter on short periods of time [1]| organisms appearing sporadically in groundater systems, called accidentals by some authors [2,3].

Stygozone

*Specialized definitions:* each of the different ecological zones in which the subterranean waters can be divided [8].

Substygophile

*Specialized definitions:* aquatic subtroglophile (author's modification).

* Syckerquelle

*Specialized definitions:* equivalent to "helocrene" spring, in German.

Thalassoid

*Specialized definitions:* stygobiont, even living in continental groundwaters, is related to marine groups and derived from marine ancestor and not closely related to surface freshwater fauna [2].

Thalassopsammal

*Specialized definitions:* biotope in the sand interstitial saline groundwaters [8].

Thalassopsammon

*Specialized definitions:* ecosystem in the sand interstitial saline groundwaters [8].

Thalassopsephal

*Specialized definitions:* biotope in the gravel interstitial saline groundwaters [8].

Thalassopsephon

*Specialized definitions:* ecosystem in the gravel interstitial saline groundwaters [8].

Thalassostygobiont

*Specialized definitions:* stygobiont living in salt groundwater [2].

Thalassostygophile

*Specialized definitions:* stygophile living in saline groundwater (author's modification).

Thalassostygoxene

*Specialized definitions:* stygoxene living in saline groundwater (author's modification).

Torrenticole (*derivations:* torrenticolous)

*Merriam & Webster:* an organism that lives in swiftly flowing water [6].

*Specialized definitions:* fauna living in flowing water [6].

Troglostygal

*Specialized definitions:* the biotope in the sand interstitial waters associated to cave rivers [8].

Troglostygon

*Specialized definitions:* the ecosystem in the sand interstitial waters associated to cave rivers [8].

Troglostygopsammal

*Specialized definitions:* biotope in the sand interstitial waters associated to cave rivers [8].

Troglostygopsammon

*Specialized definitions:* biocoenosis in the sand interstitial waters associated to cave rivers [8].

Troglostygopsephon

*Specialized definitions:* biotope in the gravel interstitial waters associated to cave rivers [8].

Troglostygopsephon

*Specialized definitions:* biocoenosis in the gravel interstitial waters associated to cave rivers [8].

#### 1.2.4. Jargon relative to superficial subterranean habitats

Alluvial MSS

*Specialized definitions:* streambeds of temporary watercourses as the result of the accumulation of rocky fragments, pebbles and gravel, deposited in situ by periodic flooding [25,26].

Bedrock MSS

*Specialized definitions:* MSS-like habitat forming due to the weathering of the bedrock at the interface between soil and parent rock, covered over time by an evolving soil [25].

Colluvial MSS (*derivations:* bare colluvial MSS, Ice-bearing taluses)

*Specialized definitions:* fragments of rocks accumulating at the bottom of rocky walls, covered over time by an evolving soil [25].

Hyporheic refuge hypothesis

*Specialized definitions:* hypothesis that the hyporheic could serve as a refuge for the invertebrate inhabitants of surface streams from environmental stresses, including flooding and drying [3].

Macrocaverns

*Specialized definitions:* underground voids >20 cm, especially caves and lava tubes [3].

Mesocaverns

*Specialized definitions:* cavities smaller than caves, between 0.1 and 20 cm in diameter [2–4].

Mesocavernous shallow substrata (*derivations:* mesocavernous superficial strata, mesocavernous superficial substra, mesocavernous shallow strata, mesocavernous rock strata)

*Specialized definitions:* equivalent to *Milieu souterrain superficiel* [25].

Mesovoids

*Specialized definitions:* equivalent to mesocaverns [25].

Mesovoid superficial substratum (*derivations:* mesovoid superficial strata, mesovoid shallow substrata, mesovoid shallow strata)

*Specialized definitions:* equivalent to *Milieu souterrain superficiel* [25].

Microcavern

*Specialized definitions:* cavities smaller than mesocaverns, less than 1 mm in diameter [3].

Microvoid

*Specialized definitions:* equivalent to microcavern.

*Milieu souterrain superficiel*

*Specialized definitions:* interconnected cracks and crevices in scree slopes and similar habitats [3,4,25]| The system of empty air-filled voids within rocky fragments, usually regarded as suitable for the survival of specialized subterranean organisms [25].

MSS (*derivations:* MSSD, MSSs, MMS)

*Specialized definitions:* official acronym for *Milieu souterrain superficiel* [25].

MSS/UHZ

*Specialized definitions:* official acronym for *Milieu souterrain superficiel*, but used by scientists who want to acknowledge that the MSS was described independently by Uéno and Juberthie et al. ; equivalent to MSS [25,27,28].

MSSv

*Specialized definitions:* official acronym for volcanic MSS [25].

Shallow subterranean habitat (*derivations:* superficial subterranean habitat, superficial subterranean environment, SSH)

*Specialized definitions:* This collective term is used to describe aphotic subterranean habitats close to the surface, harboring species with morphological and physiological adaptations to subterranean life. According to the original classification, SSHs include lava tubes, leaf litter, deep soil strata, water-filled epikarst, seepage springs, hyporheic habitats, and the MSS [4] .

Shallow underground compartment (*derivations:* shallow underground environment, superficial underground environment, shallow underground space, superficial underground space, superficial subterranean enclosure, shallow subterranean enclosure, subterranean superficial media, subterranean underground compartment, superficial hypogean compartment, spelean corridor)

*Specialized definitions:* equivalent to *Milieu souterrain superficiel* [25].

SUC

*Specialized definitions:* official acronym for superficial/shallow underground compartment; equivalent to MSS [25].

Terrestrial interstitial habitat

*Specialized definitions:* equivalent to MSS [25].

UHZ

*Specialized definitions:* official acronym for upper hypogean zone; equivalent to MSS [25].

Upper hypogean zone (*derivations:* upper hypogean habitat, upper shallow compartment, upper superficial compartment)

*Specialized definitions:* equivalent to *Milieu souterrain superficiel* [25].

Volcanic MSS

*Specialized definitions:* volcanic materials accumulating on the substrate, forming a network of voids, microspaces and fissures which are covered over time by an evolving soil [25].

#### 1.3. Jargon related to sampling methods and miscellaneous jargon

Baermann funnel

*Specialized definitions:* modification of the Berlese funnel, used to force nematodes out of the soil by filing the funnel with warm water, driving the nematodes into a vessel below [3].

Berlese funnel

*Specialized definitions:* a device to extract invertebrates from soil and leaf litter [3].

Biospeleology

*Merriam & Webster:* the biological study of cave-dwelling organisms

*Specialized definitions*: the study of life in caves and other underground environments except for interstitial ones and the meiofauna [5] | the science that studies living organisms in caves, their origin, phylogeny, adaptations, ecology, distribution, etc [1].

Bou-rouch pump

*Specialized definitions:* a special hand pump designed to collect water samples from shallow interstitial aquifers [3,4].

Karaman-chappuis method

*Specialized definitions:* a method used to collect interstitial fauna consisting of digging a hole 1-1-5 meters deep is dug in the banks of a river. Ground water and the fauna it contains rapidly filter through the sand and gravel and fill the hole. The organisms present can be simply removed with a net [6].

Meiobenthology

*Specialized definitions:* study of the meiobenthos [21].

Phreatobiological net (*derivation*: Cvetkov net)

*Specialized definitions:* a type of vertical net design to collect samples in wells and boreholes [29,6].

Phreatobiology

*Specialized definitions:* study of biology of the groundwater [3,4,22].

Racovitzan impediment

*Specialized definitions:* We cannot describe, map, analyze and conserve the biodiversity of the environments we never explored and mapped [30].

Speleobacteriology

*Specialized definitions:* study of cave microbial life [6].

Speleobiology

*Specialized definitions:* branch of biology dealing with subterranean organisms and their habitats [3,4].

Speleocommunity

*Specialized definitions:* the community of scientists working in caves; see also speleobiology.

Speleodiver *(derivations:* speleodiving, speleosub)

*Specialized definitions:* cave diver.

Speleogenesis

*Specialized definitions:* the origin and development of caves [5] | the process of cave formation [1] | the process of the origin of solution caves, and the branch of knowledge about it. In a wider sense this term means not only the very origin but also the entire life history of caves from gestation to obliteration (complete filling or decay) [2].

Speleogenetics

*Specialized definitions:* biospeleological investigations at the genetic level [31].

Speleogenomics

*Specialized definitions:* biospeleological investigations at the genomic level [31].

Speleology

*Merriam & Webster:* the scientific study or exploration of caves

*Specialized definitions:* the scientific study of caves [5].

Stygobiology

*Specialized definitions:* study of the subterranean aquatic animals [23].

SSD

*Specialized definitions:* official acronym for subterranean sampling device [25].

Subterranean sampling device

*Specialized definitions:* any device to sample the fauna in MSS habitats [25].
